# Interpretable multi-omics machine learning reveals drought-driven shifts in plant–microbe interactions

**DOI:** 10.1101/2025.08.13.670005

**Authors:** Hayato Yoshioka, Pavla Debeljak, Soizic Prado, Yushiro Fuji, Yasunori Ichihashi, Hiroyoshi Iwata

## Abstract

**Background:** Plant-microbe interactions in the rhizosphere are central to plant growth, nutrient acquisition, and stress resilience. Although multi-omics approaches enable comprehensive profiling of different biological layers, integrating these data to understand the mechanisms underlying plant-microbe symbiosis, particularly under drought stress, remains a challenge.

**Results:** Genomic, metabolomic, and microbiome data from 198 soybean accessions grown under both control and drought conditions were integrated to identify environment-specific predictive features of the plant phenotypes. We compared best linear unbiased prediction (BLUP), genome-wide association study (GWAS), and a nonlinear machine learning model to evaluate their ability to detect informative features. The machine learning models provided flexible variable selection and outperformed linear models in capturing nonlinear dependencies. Model interpretation using SHapley Additive exPlanations (SHAP) indicated that the isoflavone derivative, daidzin, and the drought-tolerant *Candidatus Nitrosocosmicus*, were major contributors to phenotypic variation, specifically under drought stress. SHAP-based interaction networks indicated cross-omics links, including connections between daidzin, gamma-aminobutyric acid (GABA), and *Paenibacillus*.

**Conclusion:** The proposed interpretable machine learning approach for plant phenotype prediction identified multi-omics biomarkers and interactions, providing insights into plant adaptation to drought stress through environment-dependent rhizosphere networks and symbiotic associations.

## Background

Interactions between plants and microbes are pivotal in regulating plant growth (Ulrich et al., 2019; Xiong et al., 2020). In particular, rhizosphere microorganisms are crucial for plant health as they enhance nutrient uptake and strengthen plant resilience, especially under abiotic stresses, such as drought (Kristin and Miranda, 2013; Alemu et al., 2024). Therefore, understanding the rhizosphere microbiome is essential for deciphering the mechanisms underlying plant-microorganism symbiosis, particularly those related to drought tolerance.

Recent advances in multi-omics technologies have enabled the concurrent profiling of genomic (Kajiya-Kanegae et al., 2021), metabolomic (Guo et al., 2023), and microbiome data (Marco et al., 2022). These inter-omics datasets provide new opportunities to unravel the complex relationships among various biological systems. However, the integration and interpretation of such diverse data types pose significant computational and statistical challenges.

Best Linear Unbiased Prediction (BLUP) is a widely used linear mixed model in plant and animal breeding (Meuwissen et al., 2001; VanRaden, 2008). It leverages genome-wide markers to predict phenotypic traits by capturing linear additive genetic contributions. While BLUP can incorporate multiple omics layers through multi-kernel approaches, it remains limited to modeling linear relationships among features.

Genome-wide association studies (GWAS) are popular for identifying individual SNP markers associated with phenotypic variation (Atwell et al., 2010; Ott et al., 2015). However, GWAS typically analyzes genome-wide markers in isolation. When evaluating the significance of the interactions between genome-wide markers and other omics variables, the number of possible combinations and multiple testing burden may increase exponentially, making such analyses infeasible.

In contrast, machine learning approaches, including Random Forest (RF; Breiman 2001), are capable of nonlinear modeling and feature selection in multi-omics contexts. RF constructs a collection of decision trees and aggregates their predictions, providing both high predictive performance and feature importance scores for biological features (Acharjee et al., 2016; Pfeifer et al., 2022). Nonlinear dependencies across omics layers have been observed in integrative studies (Zhao et al., 2022; Shen et al., 2024). Recently, our group demonstrated that the RF model can effectively capture such complexity and outperform traditional linear models, such as BLUP, in predictive tasks when nonlinearity exists in multi-omics interactions (Yoshioka et al., 2025).

SHapley Additive exPlanations (SHAP) is a game-theoretic approach that quantifies the marginal contribution of each feature by comparing model predictions across all possible subsets of input features (Lundberg and Lee, 2017). Unlike traditional feature importance scores, SHAP values are both model-specific and globally consistent, offering insights into the interactions among features and their influence on predictions (Lundberg et al., 2019).

Recent studies have employed RF and SHAP to interpret phenotype prediction models, including those used for flowering time (Wang et al., 2024). In addition, SHAP has been used with microbiome data to identify features distinguishing between drought and control conditions (Hagen et al., 2024).

However, to date, no study has systematically integrated metabolomic, microbiomic, and genomic data to identify interacting biomarkers that bridge plant–microbe interactions. Moreover, the application of SHAP-based interpretability to uncover such cross-domain interactions in plant–microbe multi-omics datasets remains unexplored.

Due to the high-dimensional nature of the interaction analysis, we employed XGBoost (Chen and Guestrin, 2016), an ensemble tree-based gradient boosting method. Unlike RF, which aggregates independently grown trees, the gradient boosting model constructs trees sequentially by optimising a loss function, enabling efficient computation in high-dimensional settings.

In this study, we specifically focused on drought-induced shifts in plant-microbe interactions. By comparing plants grown under drought and control conditions, we aimed to capture how the plant-microbe-metabolome network collectively responds to water deficit stress.

Our analysis was conducted in three steps. First, we compared the BLUP and RF models to assess their respective abilities and identify biologically meaningful features from multi-omics datasets. Second, we employed SHAP to elucidate condition-specific features and cross-omics interactions that drive phenotypic variation. Finally, through this integrative multi-omics framework, we identify drought-responsive biomarkers and uncover condition-specific regulatory mechanisms that underpin adaptive responses in plant-soil microbiome systems.

## Methods

### Soybean Multi-Omics Data

We analysed multi-omics datasets collected from a common panel of *N* = 198 soybean accessions (Fig. 1). All omics and phenotype measurements were obtained from field trials conducted in 2019 under two watering conditions: watering (control) and non-watering (drought) to assess the effects of drought stress (Toda et al., 2022; Sakurai et al., 2022). The datasets included the following:

- **Whole-genome SNP genotypes (genome)**: a 198 × 425,858 matrix, where each column corresponds to a single nucleotide polymorphism (SNP) marker in the soybean genome (Kajiya-Kanegae et al., 2021).
- **Rhizosphere metabolome**: a 198 × 263 matrix of metabolome features obtained from rhizosphere samples. Each column represents the normalized peak area of a distinct metabolome feature detected by tandem mass spectrometry (Dang et al., 2025).
- **Rhizosphere microbiota profiles (microbiome)**: a 198 × 16,457 matrix generated by 16S rRNA gene amplicon sequencing of DNA extracted from rhizosphere samples. Each column represents the relative abundance of a bacterial taxon (Dang et al., 2025).
- **Plant phenotype**: a 198 × 9 matrix. These traits include biomass-related features such as shoot and leaf fresh/dry weights, growth stage, plant height, main stem length, number of nodes, and number of branches.

**Fig. 1.**
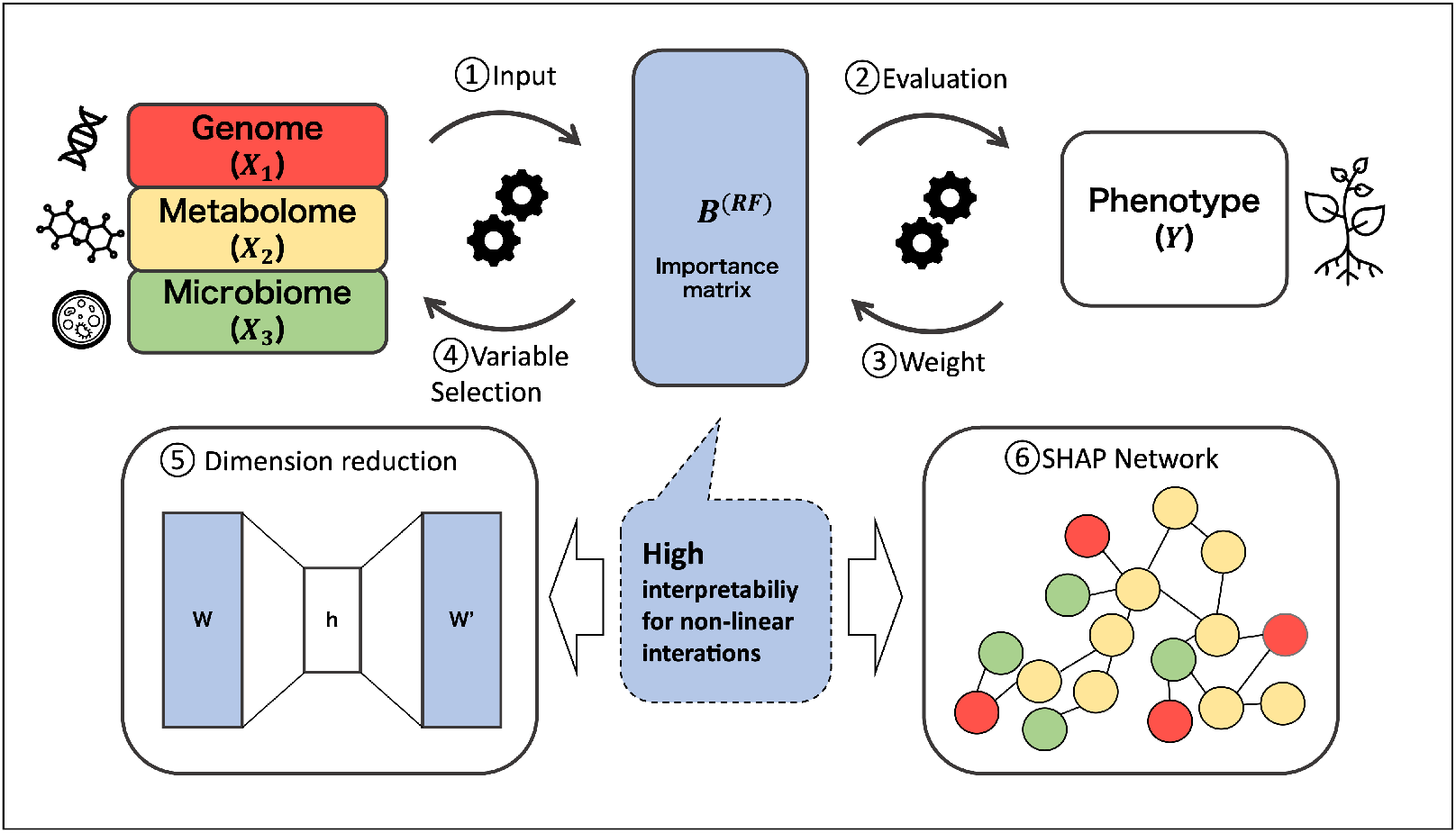
Concept of our framework. (1) Multi-omics data (Genome, Metabolome, Microbiome) are integrated as a single matrix and introduced into a soybean phenotype prediction model. (2) Phenotype (Y) is predicted using RF. (3) The model is analyzed retrospectively from the output Y to optimize the weights (importance: *B*^(RF)^) of predictive factors within the omics data. (4) Variable selection is performed based on the nonlinear matrix *B*^(RF)^, and a new prediction model is built using the selected variables. (5) Relationships among omics layers are interpreted in a low-dimensional latent space. (6) SHAP for network analysis of omics factors.

The soybean accessions used in this study were selected from the Global Soybean Minicore Collection (Kajiya-Kanegae et al., 2021). Furthermore, all rhizosphere microbiota profiles (microbiome) and rhizosphere metabolome data were collected from the same field plot as the phenotypic measurements. This design ensured consistency across samples and enabled integrative modeling across multiple omics layers.

To reduce the computational burden in the SHAP analysis of the RF model, SNPs were pruned based on linkage disequilibrium (LD). A representative subset of 3,078 SNPs (LD threshold = 0.001) was selected from an initial set of 425,858 SNPs (LD threshold = 0.95). Despite the relatively small size compared with the full data, the 3,078-SNP subset provided approximate coverage of the entire genome loci (Fig. B1). Additionally, the 3,078-SNP subset achieved predictive performance comparable to that of larger sets containing approximately 10k to 34k SNPs in phenotypic predictions (Fig. B2). The objective was to evaluate variables rather than optimise predictive performance. Thus, considering interpretability, computational limits in RF, and the need to balance the number of variables with those in the microbiome and metabolome datasets, we selected the 3,078-SNP set for subsequent RF analysis.

In the field trial, both microbiome and metabolome datasets were obtained from the same plots used for the phenotypic measurements. Note that these datasets were also recently utilized in Dang et al. (2025). Although they were collected from the same field, in the present study, the microbiome data were newly analyzed and integrated according to our experimental design, while the metabolome data were derived from those previously processed in Dang et al. (2025).

The observed phenotypic values and omics features were filtered to remove missing values. After filtering, the dataset sizes (accessions × features) were as follows. For the control condition: metabolome (182 × 255), microbiome (182 × 5424), and phenotype (182 × 9). For the drought condition: metabolome (179 × 253), microbiome (179 × 4771), and phenotype (179 × 9). All datasets were subsequently scaled prior to modeling. Details of the omics data processing for integration are provided in Appendix A.

See the supplementary information in Appendix A for details on data collection and experimental conditions. The soybean raw genome sequences are available in the DDBJ Sequence Read Archive (DRA) under the BioProject accession number PRJDB7281, as previously processed and reported by Kajiya-Kanegae et al. (2021). The microbiome raw read sequences were originally processed and are available in the DDBJ DRA under the BioProject accession number PRJDB37881. The metabolome raw data were previously processed and analyzed by Dang et al. (2025), and are available in the RIKEN DropMet database (http://prime.psc.riken.jp/menta.cgi/prime/dropindex;ID:DM0071). The plant phenotype data were originally processed and analyzed in this study. The microbiome, metabolome, and phenotype data are also provided as supplementary materials.

### Linear Mixed Model Analysis (BLUP)

To estimate the contributions of individual features to phenotypic variation, we imple-mented a linear mixed model framework based on BLUP (Meuwissen et al., 2001; VanRaden, 2008).

We have a feature matrix **X** ∈ ℝ^*n*×*p*^ and a response matrix **Y** ∈ ℝ^*n*×*q*^ . Each omics layer was used to construct a corresponding kernel matrix **K**_*l*_, which captures the pairwise similarities among samples. The univariate mixed model was performed for each phenotype *j* = 1,…,*q* as follows:

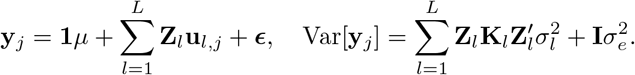

The multi-kernel linear mixed effects model was computed using the R package `RAINBOWR` (version 0.1.35) (Hamazaki and Iwata, 2020). The model was applied inde-pendently to each phenotype *j* = 1,…,*q* to obtain trait-specific random effects **u**_*l,j*_ for each omics layer *l*. Assuming that each kernel matrix was based on a linear inner product, we reconstructed the feature-level coefficients for each trait and omics layer using:

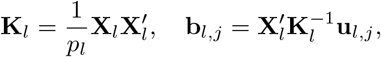

where 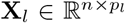 is each feature matrix (genome, metabolome, or microbiome), and 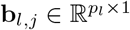 is the resulting coefficient vector.

These feature matrices were used to construct the corresponding kernel matrices **K**_*l*_ for every layer. All coefficient vectors were concatenated across the traits and layers to form a global coefficient matrix:

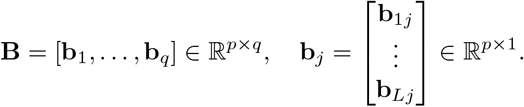

To enable comparison across traits, each column of **B** was standardised to have a mean 0 and a variance 1:

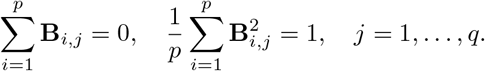

To examine the structure of the feature-trait relationships, we applied singular value decomposition (SVD) to the coefficient matrix:

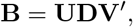

where **U** and **V** are the left and right singular vectors, respectively, and **D** is the diagonal matrix of singular values. The phenotypes and features were projected into the latent space using 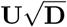 and 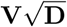, respectively.

### Nonlinear Feature Importance Using RF

We also implemented a nonlinear regression framework using RF to estimate feature contributions. The feature matrix **X** ∈ ℝ^*n*×*p*^ was constructed by concatenating multiomics layers, while the response matrix **Y** ∈ ℝ^*n*×*q*^ represents the biomass-related phenotypic traits. Details of the data preprocessing are provided in Appendix A.

For each phenotype *j* = 1,…, *q*, we trained an independent RF model using the R package `ranger` (version 0.16.0) (Wright and Ziegler, 2017). Hyperparameter tuning was performed using a grid search. See Appendix C and Fig. B3 for details. We confirmed that the predictive performance of the RF model fitted to the original data exceeded the null distribution of predictive performance obtained by refitting the model after randomly permuting the response vector (Fig. B4).

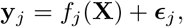

where *f*_*j*_ denotes the RF model trained for phenotype *j*. After training, the feature importance scores were extracted. Let **I** ∈ ℝ^*p*×*q*^ denote the resulting matrix of feature contributions, where **I**_*i,j*_ indicates the contribution of feature *i* to phenotype *j*.

The importance values from the RF were all positive. For the convenience of variable selection, each column of **I** ∈ ℝ^*p*×*q*^ is normalised such that the total sums to 1 for each phenotype:

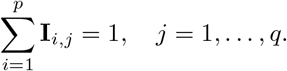

Analogous to the coefficient matrix **B** derived from BLUP, the normalised importance matrix **I** ∈ ℝ^*p*×*q*^ from the RF models can be treated as a feature-by-trait matrix:

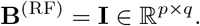

Here, each column of **B**^(RF)^ represents the relative contribution of each feature to the prediction of a specific phenotype, aggregated across the trees in the forest (Fig. 1). While the entries of **B**^(RF)^ do not correspond to regression coefficients in a strict parametric sense, this matrix serves as a comparable construct for downstream analyses, such as dimensionality reduction or visualisation of feature-trait associations. Following the BLUP, we can perform the SVD for **B**^(RF)^ to visualise the structure of feature-trait relationships.

Leaf dry weight, a commonly used indicator of plant biomass that reflects the size of the primary photosynthetic organ, showed representative predictability and consistent associations with omics features across all three omics layers (Fig. 2). Therefore, downstream GWAS and SHAP analyses were conducted using leaf dry weight as a representative phenotype for interpretability and computational feasibility.

**Fig. 2.**
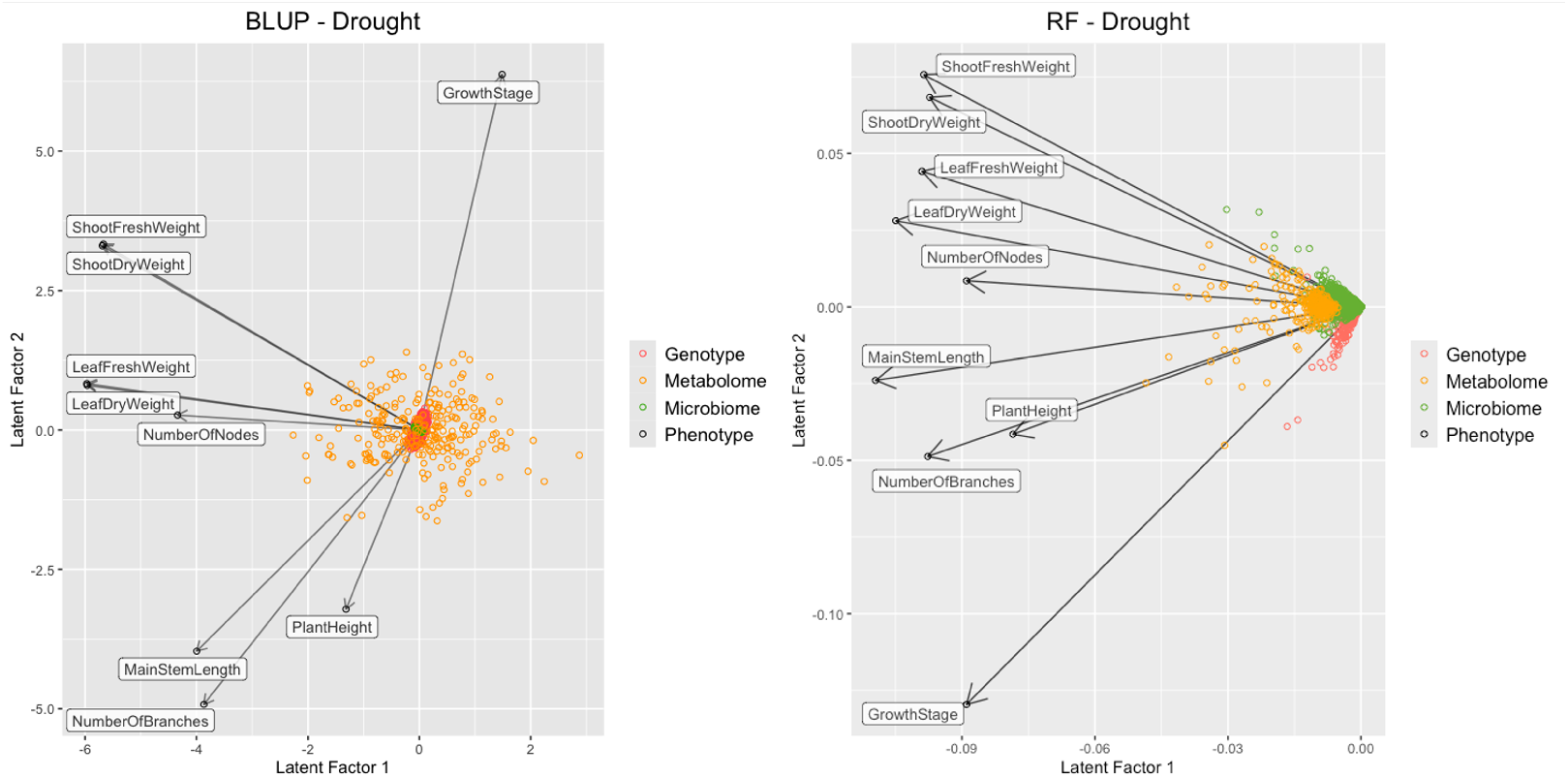
Feature–trait structures derived from BLUP and RF for predicting biomass-related phenotypes under drought conditions. Feature importance matrices from both models were subjected to singular value decomposition (SVD), and the first two latent factors are visualized. Points represent features and phenotypes projected into the latent space. Colors indicate data types (genotype, metabolome, or microbiome), and phenotype vectors are shown as arrows.

### Genome-wide association study (GWAS)

GWAS is a linear approach for identifying SNP markers associated with phenotypic variation (Atwell et al., 2010; Ott et al., 2015). Here, we employed GWAS in a conventional setting to evaluate the baseline performance and provide a comparison with SNP selection results from RF importance.

GWAS was performed using a linear mixed model (LMM) implemented in the R package `gaston`(Perdry and Dandine-Roulland, 2023) (version1.6), incorporating three principal components (PCs) derived from the eigen decomposition of the GRM to account for population structure.

The full set of SNPs (425,858 markers, LD threshold = 0.95) was filtered using a minor allele frequency (MAF) threshold of 0.05 and then used in the GWAS model. After modeling, SNPs with a significance threshold of *p<* 1 × 10^−5^ were retained for further analysis.

### SHAP: Individual-Features Attribution

To investigate the individual contributions of multi-omics features to phenotype prediction (hereafter referred to as individual feature contributions), we applied SHAP to the trained RF models. SHAP is a game-theoretic approach that quantifies the marginal contribution of each feature by comparing the model predictions across all possible subsets of the input features.

Formally, the SHAP value for feature *i* is defined as:

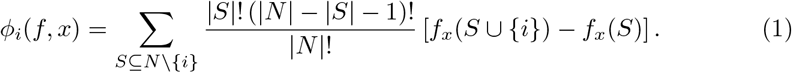

where *N* is the set of all input features, *S* is a subset that excludes feature *i*, and *f* is the prediction model. The *f*_*x*_(*S*) denotes the model prediction for input instance *x* when only the features in *S* are considered. The left-hand side of equation (1) is a combination weight designed to give equal consideration to all possible feature orders. The right-hand side represents the marginal contribution to the prediction when feature *i* is added to subset *S*. By combining these components, feature importance can be assigned fairly, making SHAP particularly suitable for elucidating the nonlinear effects of features in multi-omics data.

SHAP values were computed for each sample using the `iml` R package (version 0.11.4) (Molnar et al., 2018) and then averaged to obtain global feature importance scores.

After RF modeling under drought and control conditions separately, omics features were ranked based on their total variable importance scores, calculated for each feature *i* as follows:

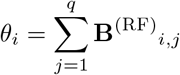

were retained, where **B**^(RF)^_*i,j*_ denotes the contribution of omics feature *i* to phenotype *j*.

Here, we selected features with a total RF importance score of *θ*_*i*_ *>* 0.01 under either drought or control conditions, yielding a total of 159 features for evaluation. This threshold was chosen to retain features that exhibited non-negligible importance, given that the distribution of the feature importance scores was zero-inflated (Fig. B5). From a practical perspective, applying such a cutoff provides a reasonable preprocessing step for SHAP analysis because it reduces the computational burden while preserving the major informative signals.

Using this subset of 159 features, the SHAP values were calculated to identify environment-dependent differences in the relative contribution of individual features. For each condition, the absolute SHAP values were first calculated for each feature across all samples and then averaged. The ratio of the mean absolute SHAP values (drought divided by control) was calculated to highlight the condition-specific features of the model.

### SHAP: Interaction-Features Attribution

#### Condition Specific Interactions

Although SHAP individual-feature attribution values quantify each omics feature, they do not capture interaction effects. To address this, we applied SHAP interaction analysis to characterise the pairwise relationships among the features (Molnar, 2025).

SHAP interaction values decompose the prediction into individual effects and pairwise interactions.

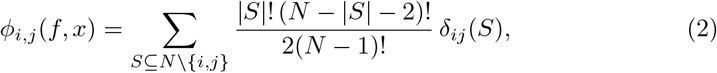

where *i* ≠ *j* and

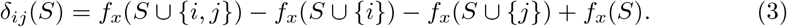

The weighting factor in equation (2) ensures the symmetry consideration of all feature orderings, whereas the term *δ*_*ij*_(*S*) isolates the second-order marginal contribution in equation (3).

Although some features exhibited only minor individual effects on the phenotype, they might contribute to phenotypic variation through cooperative or context-dependent interactions with other omics layers. To capture such synergy-driven patterns, we adopted a broader feature set (*θ*_*i*_ *>* 0.001 in both conditions), which facilitated the identification of interaction effects that were not reflected in the individual feature importance. The distribution of feature importance values is shown in Fig. B5.

To evaluate the sensitivity of the inferred interaction network to the RF importance threshold *θ*, we conducted a robustness analysis by varying *θ* from 0.001 to 0.01. SHAP interaction scores were first computed under the most permissive setting (*θ* = 0.001). For each stricter threshold, nodes retained according to RF importance were identified for each condition, and the interaction network was restricted to edges whose two incident nodes remained selected.

This yielded a total of 2,729 features for the evaluation. Based on these features, we constructed omics feature interaction networks, where nodes represent individual features and edges denote the strength of interaction between feature pairs.

In high-dimensional SHAP interaction analysis, the number of feature pairs increases exponentially with the number of input variables. Due to this computational burden, the interaction values were computed using the `shap.TreeExplainer` from the `shap` Python package applied to an XGBoost-based gradient boosting model (Chen and Guestrin, 2016), as RF-based implementations were computationally prohibitive in this setting. Since both RF and gradient boosting are tree-based ensemble methods, the resulting SHAP interaction values captured the same class of nonlinear feature interactions underlying the nonlinear prediction models.

Hyperparameter tuning was conducted so that the same set of tuned hyper-parameters could be applied to both the control and drought conditions, ensuring comparability between environmental settings. See Appendix C and Fig. B3 for implementation details.

SHAP interaction values were calculated for each sample and averaged to obtain the global 2,729 ≠ 2,729 symmetrical interaction matrices for each condition. To highlight the environment-specific interactions, we computed a differential SHAP interaction matrix as:

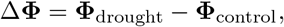

where **Φ** denotes a SHAP-based pairwise interaction matrix. Positive values of Δ**Φ** indicated stronger interactions under drought stress, whereas negative values indicated interactions that were more pronounced under control conditions.

The two-step SHAP analysis—first focusing on individual feature importance, then exploring pairwise interaction features—provides a comprehensive and interpretable view of how genomic, metabolomic, and microbial features jointly influence plant phenotypes under contrasting environmental conditions.

#### Null-Distribution-Based Significance Assessment

To assess the statistical significance of each differential interaction, a permutation-based null distribution was constructed. For each condition, the response vector was randomly permuted. The XGBoost model was then refitted to the permuted data, and SHAP interaction matrices were computed. For each permutation replicate *b* ∈ {1,…, 100}, a null differential interaction matrix was calculated as:

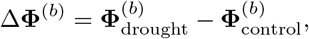

representing the distribution of interaction differences expected under the null hypothesis of no association between predictors and observed values (phenotypes).

All off-diagonal (upper-triangular) entries from the *B* = 100 null differential interaction matrices were concatenated to form a pooled null distribution, from which the mean *μ*_0_ and standard deviation *σ*_0_ were estimated. The observed differential interaction values were converted into Z-scores as:

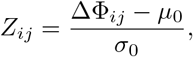

Here, edges with large absolute Z-scores were interpreted as highly environment-specific SHAP interactions.

Edgewise empirical *p*-values were computed for each feature pair (*i, j*) using two-sided tests based on the null replicates (Phipson and Smyth, 2016).

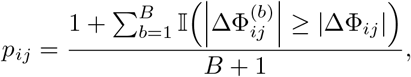

To examine the robustness of the inferred interaction network with respect to the choice of the Z-score threshold, we systematically constructed networks using a range of alternative cutoffs ( |*Z*| ≥ 10, 20, 30, 40, 50, and 60). For each threshold, the cor-responding two-sided empirical *p*-values were jointly evaluated to ensure consistency between effect size and statistical extremeness.

Although the overall qualitative network structure was preserved across the thresholds, a cutoff of |*Z*| ≥ 30 was selected for the downstream analyses as a pragmatic compromise between statistical stringency and network interpretability. This threshold corresponds to the extreme tail of the pooled null distribution of SHAP interaction differences (Fig. B6) and effectively filters out interactions that are likely attributable to resampling noise. Under this cutoff, the resulting network comprised 160 connections, representing only 0.004% of all possible feature pairs, yielding a sparse yet interpretable representation of robust environment-specific interactions.

#### Stability Analysis of Interaction Edges using Bootstrap Resampling

Because SHAP interaction matrices are estimated from finite samples, the individual edge differences in Δ**Φ** may be sensitive to resampling variability. To assess the robustness of the differential interactions, edge-level stability was quantified using an additional bootstrap procedure applied to the observed data.

Specifically, for each condition, we generated 100 bootstrap replicates by resampling individuals with replacements and refitting the XGBoost model on the resampled pairs {(**x**_*k*_, *y*_*k*_)} . For each replicate *b*, we computed the bootstrap differential interaction matrix

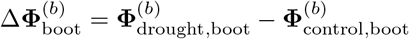

Using these replicates, the complementary stability measures for each edge were defined, (*i, j*). The presence stability was defined as the number of bootstrap replicates for which a given edge (*i, j*) is observed as nonzero.

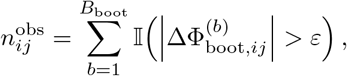

where *ε* = 10^−12^ is the small numerical tolerance. This measure aims to evaluate the reproducibility of the interaction under resampling, which is particularly relevant for sparse or zero-inflated interaction matrices.

To suppress the spurious edges driven by resampling noise, a stability-based filtering criterion was applied, and edges with 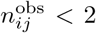 were excluded from downstream network analyses. Given the extreme sparsity of SHAP interaction matrices (≈ 0.015%, corresponding to approximately 500 of 3.7 million possible pairs), the appearance of a purely spurious edge in each bootstrap replicate could be treated as a rare event. Under a simple binomial model *X* ∼ Binomial(*B*_boot_ = 100, *p*) with *p* ≈ 1.5 × 10^−4^, the probability that the edge observed in at least two bootstrap replicates is approximately 1.1 × 10^−4^. See Appendix C.3 for further statistical justification for this conservative threshold. Thus, the stability metric: 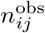 is reported alongside the Z-scores and empirical *p*-values to facilitate interpretation of the edge-level robustness.

## Results

### RF and BLUP Models

Fig. 2 presents the SVD of the coefficient matrix of BLUP and the importance matrix of RF for the drought conditions. The results demonstrate that the RF model exhibited a more pronounced and variable pattern of feature importance than the BLUP model. Notably, in BLUP, metabolites are the features most correlated with phenotypes, while other features (such as the genome and microbiome) exhibit a shrinkage pattern, indicating that metabolome features primarily contribute to phenotypic prediction. Under control conditions, a similar trend was observed (Fig. B7).

### SHAP analysis of individual-features

SHAP analysis suggests key condition-specific features contributing to phenotypic variations under drought and control conditions. Since metabolomic and microbiomic profiling do not cover the entirety of omics information, the highlighted features should be understood as those selected from the analysed subset described in the Materials and Methods.

Under drought stress, the most highly contributing features were predominantly flavonoids, including isoflavones (e.g. daidzin and its derivatives and genistin) and flavone/flavanone glucosides (e.g. apigenin-7-O-glucoside and naringenin-7-O-glucoside). Flavonoids play a major role in plant responses to drought stress and are associated with plant-microbe interactions (Gargallo-Garriga et al., 2018; He et al., 2022). Additionally, we identified amino acids and their derivatives (asparagine, ornithine, and indole-3-carboxyaldehyde) as major contributing features.

Furthermore, *Candidatus Nitrosocosmicus*, a clade of ammonia-oxidising archaea (AOA), was particularly selected under drought. This clade is notable for its ability to produce manganese catalase (MnKat), which decomposes hydrogen peroxide (H_2_O_2_), a major reactive oxygen species (ROS) (Sauder et al., 2017). This enzymatic capacity enables *Ca. Nitrosocosmicus* to survive in oxidative environments, such as drought-stressed soils. Its dominance under ROS stress conditions highlights its potential role in AOA-plant interactions (Lee et al., 2024), consistent with our findings.

In contrast, under control conditions, genetic markers contributed significantly to the observed traits. SHAP re-identified the SNP `Chr06-15056895` as an important feature, which was also detected by GWAS (Fig. B8). This SNP encodes a copper amine oxidase family protein involved in amine metabolism, tissue differentiation, and organ development in plants (Tavladoraki et al., 2016). An SNP `Chr04-22035062`, located near the gene *Glyma*.*04G140400*, was also selected. This gene encodes an oxidoreductase enzyme that is potentially involved in plant growth (Garrone et al., 2015). Another notable marker, SNP `Chr04-16278976`, is located near *Glyma*.*04G127100* (Valliyodan et al., 2019). This gene encodes a receptor-like kinase that contains both a protein kinase-like domain (IPR011009) and a Gnk2-homologous domain (IPR002902) and is involved in signal transduction pathways (Shiu and Bleecker, 2001).

In addition to genetic factors, *Duganella* was identified as an important microbial contributor under control conditions in this study. Several species of *Duganella* are known for their plant growth-promoting properties, including the solubilisation of essential nutrients, such as phosphorus, potassium, and zinc (Verma et al., 2014).

Some metabolites, such as gamma-aminobutyric acid (GABA), were selected in both control and drought conditions; however, they were not classified as condition-specific factors because their levels were similarly elevated under both conditions. Further details of each condition are presented in Fig. B9.

### SHAP analysis of interaction-features

SHAP interaction values enabled a detailed investigation of multi-omics feature interactions. Condition-specific connectivity patterns were identified by comparing network differences between the drought and control conditions using a Z-score threshold of |*Z*| ≥ 30 (Fig. B6). Under this threshold, major features were identified based on their connectivity with other features, including GABA and daidzin, both of which are known to promote plant growth and to influence rhizosphere microbiome composition, particularly under drought stress (Bennett et al., 2004; Xu et al., 2021; Ramesh et al., 2015; Gargallo-Garriga et al., 2018; Zhao et al., 2025). The corresponding networks were obtained by using alternative Z-score thresholds ( |*Z*| ≥ 10, 20, 30, 40, 50, 60), and are shown in Fig. B10.

Several of these interactions correspond to the known biological relationships. Notably, the model identified a strong interaction between daidzin and the bacterium *Paenibacillus* under drought conditions. This may be supported by a previous study showing that certain strains of *Paenibacillus* produce *β*-glucosidases that catalyse the hydrolysis of glycosidic bonds in isoflavone glucosides, such as daidzin, thereby converting them into their active aglycone forms, such as daidzein (Park et al., 2013). In addition, the model suggested a biologically plausible interaction between *Paenibacillus* and GABA under drought conditions. This aligns with reports that *Paenibacillus* can express glutamate decarboxylase, facilitating the conversion of glutamate to GABA (Singh et al., 2023).

Under control conditions, a notable interaction was observed involving SNP `Chr04-22035062`, which was also selected in the individual-feature analysis, located downstream of the gene *Glyma*.*04G140400* (oxidoreductase enzyme), potentially involved in metabolic responses to GABA.

We additionally evaluated the sensitivity of the inferred interaction network to the RF importance threshold *θ*. The main biological interactions reported in the network remained stable across a wide range of *θ* values (Table B1). For example, the interaction between *Paenibacillus* and daidzin was retained for 0.001 ≤ *θ* ≤ 0.003, the interaction between *Paenibacillus* and GABA for 0.001 ≤ *θ* ≤ 0.006, the interaction between `Chr04_22035062` and GABA for 0.001 ≤ *θ* ≤ 0.01, and the interaction between *Rhodanobacter* and daidzin for 0.001 ≤ *θ* ≤ 0.005. These results confirm that the principal interaction structure was robust to moderate changes in the preprocessing threshold (Fig. B11).

To assess the robustness of the detected interactions, their stability across 100 bootstrap replicates was evaluated (Tables B2 and B3). Among bootstrap replicates, several edges showed moderate stability. The edge between SNP `Chr04-22035062` and GABA exhibited relatively higher stability (observed in 14 bootstrap replicates). The interactions between *Paenibacillus* and daidzin (6 replicates) and between *Paenibacillus* and GABA (3 replicates) appeared across resampling iterations. Similarly, an interaction between *Rhodanobacter* and daidzin was detected in 6 replicates, suggesting a reproducible association between the taxon and isoflavone-related metabolites. However, the magnitude of SHAP interaction differences (Drought − Control) was not necessarily proportional to the replicative stability (Fig. B12).

## Discussion

The SVD of the feature-trait matrices for BLUP and RF showed a more dispersed and interpretable structure for the RF-based models, particularly under drought conditions (Fig. 2 and B7). In contrast, BLUP showed more shrinking effects, which is typical when the explanatory matrix contains many variables. This suggests that a nonlinear model may be better suited for feature selection in complex biological systems, where additive linear assumptions do not hold.

The overlap between SNPs identified by GWAS and those selected by SHAP highlights the potential synergy between traditional and machine learning-based approaches (Fig. B8). For instance, SNP `Chr06-15056895`, located near the gene *Glyma*.*06G178400*, was detected by both SHAP and GWAS, reinforcing its putative role in growth-related traits. Such convergence enhances confidence in candidate biomarkers.

Our findings suggest a shift in the dominant omics layers under drought stress, highlighting the enhanced roles of metabolomic and microbial features (Fig. 3). Notably, the strong contribution of flavonoids, including daidzin, supports previous evidence that soybean roots adjust flavonoid exudation to shape the rhizosphere community in response to drought stress (Gargallo-Garriga et al., 2018; He et al., 2022). The identification of amino acids and their derivatives is also consistent with previous findings, as amino acid concentrations increase under drought conditions (Postles et al., 2016). Additionally, the selection of *Candidatus Nitrosocosmicus*, a major contributor to phenotypic variation under drought stress, further validates the SHAP model’s identification of key features in the microbial community (Sauder et al., 2017; Lee et al., 2024).

**Fig. 3.**
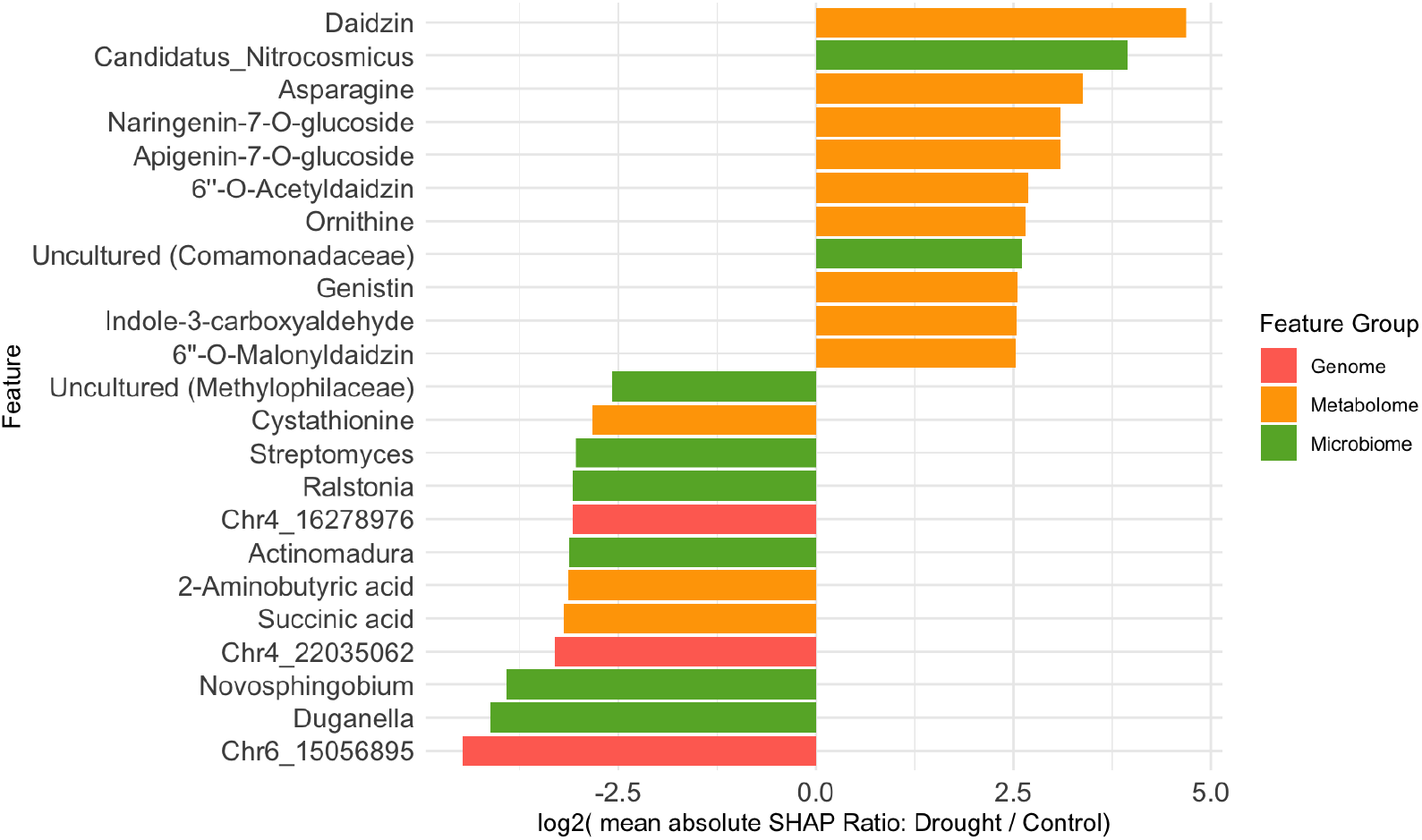
SHAP-based individual feature importance. For each control and drought condition, absolute SHAP values were computed and their ratios were calculated. The bar plot shows the log_2_ ratio of the mean absolute SHAP values (drought divided by control). Features with log_2_(mean absolute SHAP ratio) *>* 2.5 are displayed. Positive values indicate features more specific to drought conditions, whereas negative values indicate features more specific to control conditions.

Interaction analysis provided further plausible plant-metabolic-microbial links (Fig. 4). For example, daidzin, which was selected as an individual feature, was also selected as a feature of the pairwise interactions. Especially, the co-selection of daidzin and *Paenibacillus* aligns with reports that *Paenibacillus* strains produce *β*-glucosidases to hydrolyse isoflavone glucosides such as daidzin (Park et al., 2013). This enzymatic synergy may be related to ROS management. Additionally, the inferred interaction between *Paenibacillus* and GABA agrees with evidence that this bacterium encodes glutamate decarboxylase (Singh et al., 2023), which could support plant stress resilience. Interestingly, although GABA was not selected as a condition-specific feature based on individual SHAP contributions, network-level analysis revealed a strong association with drought-specific activity. This highlights the advantage of network approaches in capturing complex indirect relationships that are not apparent from individual feature contributions alone.

**Fig. 4.**
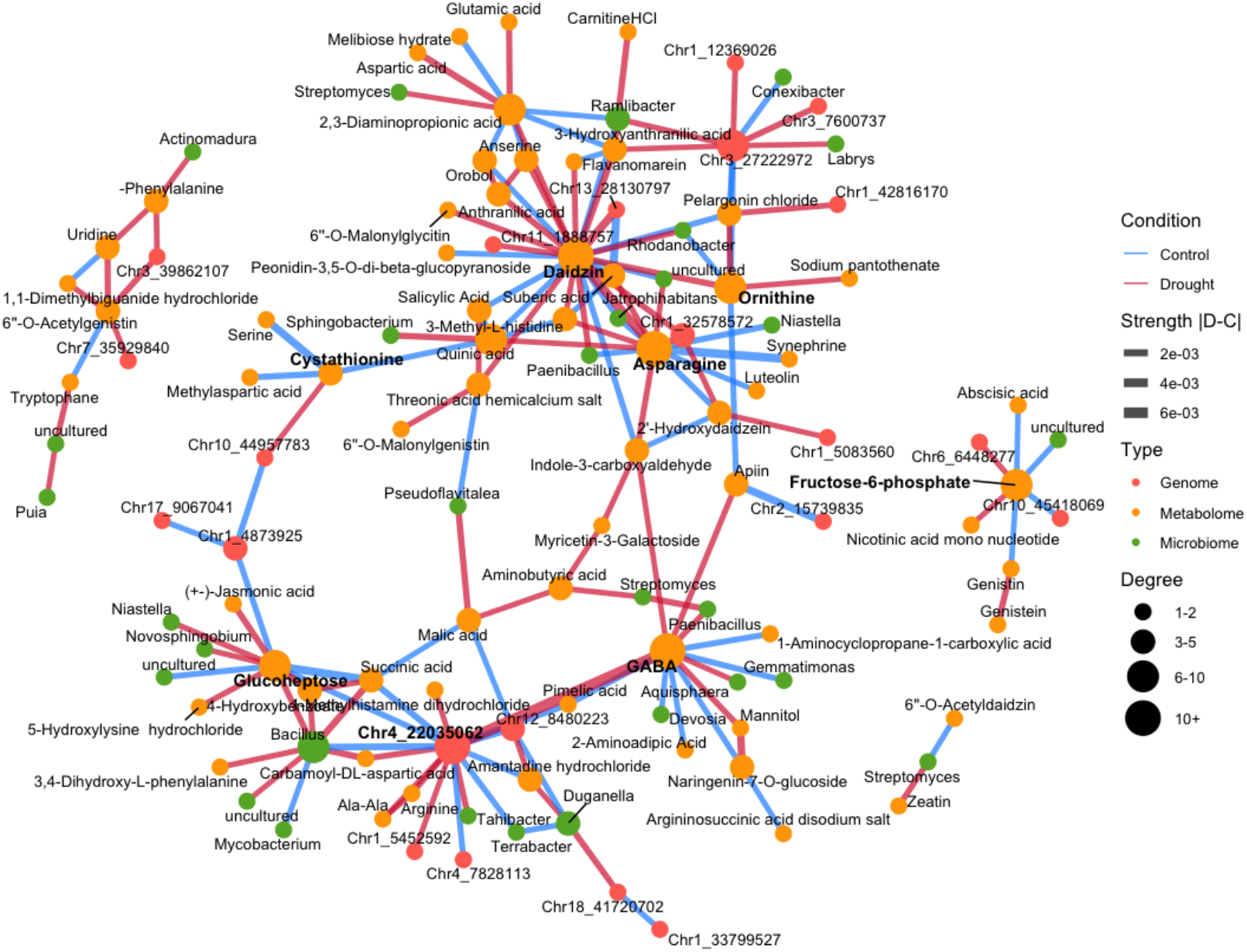
Condition-specific SHAP interaction difference network. The model was trained separately to predict biomass (leaf dry weight) under drought and control conditions. Environment-specific interactions were identified from the differential SHAP interaction matrix (drought minus control) using Z-scores derived from a pooled null distribution. Only interactions with large absolute Z-scores ( |*Z*| ≥ 30) were retained. Red edges represent interactions that are stronger under drought conditions, whereas blue edges indicate interactions that are more pronounced under control conditions. Node colors denote omics layers: genome SNP markers (red), microbiome features (green), and metabolome features (orange). Node size reflects the number of connections (degree), edge width corresponds to the magnitude of the interaction difference between drought and control, and highly connected features (hubs) are highlighted in bold font.

In addition to these known interactions, the model also suggests other potential connections between plant secondary metabolites and the microbiome that have not yet been directly reported. For example, daidzin is associated with bacteria such as *Rhodanobacter* (Singh et al., 2023), which is known to promote plant growth. GABA was also connected with *Devosia*, a plant growth–promoting genus (Zhang et al., 2024), and *Aquisphaera*, which is frequently detected in the plant rhizosphere (Thompson et al., 2025). These findings highlight the potential to elucidate previously unknown plant-microbe interactions, in which plants may use flavonoids and other metabolome features to modulate the composition of rhizosphere bacteria (Hassan and Mathesius, 2012; Gargallo-Garriga et al., 2018; He et al., 2022).

Interestingly, our analysis showed contrasting patterns of feature associations under drought and control conditions. Under drought stress, metabolomic features were particularly prominent in the modeling, suggesting scenarios in which multiple metabolites are simultaneously produced by plants to cope with drought stress. For instance, the correlation between ornithine and asparagine may reflect amino acid accumulation patterns that influence plant stress responses (Postles et al., 2016). This highlights the importance of metabolite co-regulation and network-level interactions during abiotic stress.

In contrast, under control (non-drought) conditions, genetic determinants were more strongly associated with trait expression, reflecting the predominance of intrinsic growth programs and host-regulated physiological processes in the absence of stress, that is, genetic effects outweighing environmental influences. This is supported by the identification of genotypic markers near genes involved in signal transduction, amine metabolism, and development.

Interaction edges were observed in only a limited number of bootstrap replicates; however, this pattern is a natural consequence of the extremely high-dimensional and sparse interaction spaces in which only 0.015% of all possible feature pairs exhibited non-zero SHAP interactions. A simple binomial argument indicates that observing the same interaction in as few as 2–4 of 100 bootstrap replicates is highly unlikely under a null model driven purely by sparsity. Given the empirical nonzero rate of SHAP interactions (*p* ≈ 1.5 × 10^−4^), the probability of observing an edge at least twice, three times, or four times by chance alone is of the order of 10^−4^, 10^−7^, and 10^−9^, respectively. Thus, even modest levels of replicative stability can provide non-trivial statistical support in this setting (Appendix C.3).

Some interactions exhibit large point-estimate differences but limited bootstrap recurrence, whereas others show moderate differences with higher stability (Fig. B12). This pattern may partly arise from the severe multicollinearity inherent in high-dimensional data such as those analyzed in this study, which complicates feature selection. In such settings, the model may alternate between correlated features across bootstrap samples, leading to reduced apparent stability. This issue can potentially be mitigated by incorporating stronger regularization or dimension-reduction approaches.

In the present study, we intentionally prioritized the detection and biological interpretation of interaction effects, using stability as a supportive criterion. Alternative analytical designs are also feasible. For example, greater emphasis could be placed on stability by adjusting the model complexity, regularization strength, or low-dimensional feature selection thresholds to enhance the collective recurrence of interactions. Such strategies would likely enable analyses to focus more explicitly on reproducibility.

Overall, the integrated framework enables scalable feature selection across multiple omics layers, including plant genomic data, and highlights potentially meaningful biological interactions. Despite this choice, there were some limitations. Although the proposed framework identifies synergistic associations across omics layers, it does not establish causal relationships between features. Therefore, the resulting interaction network reflects model-based associated contributions rather than direct mechanistic links, and should be interpreted as a hypothesis-generating representation to guide targeted validation.

Accordingly, further experimental studies are required to validate and extend our findings. Causal biological mechanisms can be explored through targeted experimental designs, such as quantitative trait locus (QTL) mapping and gene expression analyses, which will help refine candidate loci or genes and elucidate the underlying gene regulatory networks.

Moreover, we plan to expand this framework to incorporate larger untargeted metabolomics and microbiome datasets spanning a broader range of organisms, including fungi. Such extensions will facilitate investigations of broader ecological and ecosystem-level processes and provide deeper insights into the complex dynamics that shape these systems.

## Conclusions

This study demonstrates the utility of comparing linear and nonlinear modeling approaches to validate feature selection in multi-omics datasets. By comparing BLUP and GWAS with a nonlinear machine learning model across genomic, metabolomic, and microbiomic layers, we confirmed that the RF model captured additional nonlinear interactions, avoiding the shrinkage effects observed in linear methods. The use of SHAP further ensured that these nonlinear patterns were biologically interpretable and not model artefacts. We identified environment-specific biomarkers linked to soy-bean biomass under drought and control conditions, providing a basis for predictive breeding and microbiome-informed strategies to improve stress resilience. Future studies incorporating temporal sampling and experimental validation will help translate these multi-omics insights into practical strategies for sustainable crop production in diverse environments.

## Competing interests

The authors declare that they have no competing interests.

## Acknowledgements

We are grateful to the following researchers for their contributions to soybean data collection: Masami Hirai Yokota, Megumi Narukawa, Erika Usui, Kie Kumaishi, Takumi Satou, Shungo Kobori, Yutaka Yamada, Akito Kaga, Yusuke Toda, Kengo Sakurai, Hideki Takanashi, Yoshihiro Ohmori, Motoyuki Ishimori, Toru Fujiwara, Hirokazu Takahashi, Mikio Nakazono, Mai Tsuda, Yuji Yamasaki, Hisashi Tsujimoto, Izumi Higashida, and all the technical staff at the Arid Land Research Center, Tottori University. We appreciate the advice of Dr. Komlan Avia.

## Funding

This work was supported by JSPS KAKENHI (JP23KJ0506, JP22K21352), JST CREST (JPMJCR16O2), JST ALCA-Next (JPMJAN23D1), and JST-Mirai (JPMJMI120C7).

## Data availability

The soybean genome data were previously processed by Kajiya-Kanegae et al. (2021), and are available in the DDBJ Sequence Read Archive (DRA) under the BioProject accession number PRJDB7281.

The metabolome raw data were previously processed by Dang et al. (2025), available on the RIKEN DropMet website (http://prime.psc.riken.jp/menta.cgi/prime/dropindex;ID:DM0071). The data are also provided as supplementary materials.

The microbiome raw read sequences are available in the DDBJ DRA under the Bio-Project accession number PRJDB37881 and were originally generated and published in this study. The data are also provided as supplementary materials.

The plant phenotype data were originally processed and analyzed in this study, and the data are provided as supplementary materials.

All codes and data are available at GitHub (https://github.com/Yoska393/ShapPlantMicro).

## Contributions

H.Y., P.D., S.P., and H.I. conceived the study. Y.F. and Y.I. curated the data. H.Y. analyzed the data. H.Y., P.D., S.P., and H.I. wrote and reviewed the manuscript.

## Notes

https://github.com/Yoska393/ShapPlantMicro

### Competing Interest Statement

The authors have declared no competing interest.

### Summary of Updates

Evaluation methods, writing, authors have been corrected.

