## Appendix for "Interpretable multi-omics machine learning reveals drought-driven shifts in plant–microbe interactions"

### Appendix A Multi-omics data collection

#### A.1 Field experiment

The accessions and experimental fields were the same as those utilized by Toda et al. (2022), Sakurai et al. (2022), Dang et al. (2025). A diverse collection of 198 soybean accessions, registered with the NARO Gen-bank (https://www.gene.affrc.go.jp/), was employed. This group primarily comprises global soybean minicore collections (Kajiya-Kanegae et al., 2021). The field trial took place in 2019 at the Arid Land Research Center of Tottori University on sandy soil. Sowing was carried out at the beginning of July, and plants were thinned two weeks later. Prior to sowing, fertilizers (13, 6.0, 20, 11, and 7.0 g $m^{-2}$ of N, P, K, Mg, and Ca, respectively) were applied to the field. White mulch sheets (Dupont, Wilmington, DE, USA) were placed to minimize rain-water infiltration and to regulate soil conditions under artificial irrigation. Irrigation tubes were installed beneath the sheets to water the fields. Two watering regimes, non-watered and well-watered, were established to assess the effects of drought and control conditions. The watering treatment began after thinning and continued two weeks post-sowing, with artificial irrigation applied at a flow rate of 1.1 L $h^{-1}$ $m^{-1}$ for 5 h daily (7:00–9:00, 12:00–14:00, and 16:00–17:00). Phenotypic sampling (from the 2nd and 3rd individuals) and root sampling (from the 3rd individual) were performed at the beginning of September. These data were obtained from the same plot to facilitate the data integration for the model. The sampled root was stored at –80°C until metabolome and microbiome analyses.

#### A.2 Phenotypic Traits

The biomass-related phenotypic traits measured were dry weight of leaves (`LeafDryWeight`), dry weight of shoots (`ShootDryWeight`), fresh weight of shoots (`ShootFreshWeight`), fresh weight of leaves (`LeafFreshWeight`), growth stage (`GrowthStage`), number of nodes (`NumberOfNodes`), number of branches (`NumberOfBranches`), plant height (`PlantHeight`), and main stem length (`MainStemLength`).

#### A.3 Metabolome Analysis

Metabolome analysis of root samples was conducted as follows. Samples were freeze-dried, powdered using a mixer (Shake Master NEO BMS-M10N21, Bio Medical Science, Tokyo, Japan), and 4 mg of the powder was extracted with 1 mL of solvent (80% methanol containing 0.1% formic acid, 8.4 nM lidocaine, and 210 nM 10-camphorsulfonic acid as internal standards). The mixture was sonicated for 10 min and centrifuged at 10,000 rpm for 5 min, and the supernatant was collected as the extract. To 25 $\mu$L of the extract, 75 $\mu$L of extraction solvent was added. Twenty-five $\mu$L of this dilution was dried under $N_2$ gas, reconstituted in 250 $\mu$L of deionized water, and used as the test solution. Final concentrations of the internal standards in the test solutions were 0.84 nM for lidocaine and 21 nM for 10-camphorsulfonic acid. Following filtration through a MultiScreen 386 well-plate filter (Merck Millipore, Billerica, MA, USA), metabolome analysis was performed using a Shimadzu LCMS-8050 system coupled to a Nexera X2 UPLC system (Shimadzu, Kyoto, Japan). Chromatographic separation employed a Waters HSS T3 column (1.0 × 50 mm, 1.8 $\mu$m) at 30 °C. The mobile phases were 0.1% (v/v) formic acid in water (A) and 0.1% (v/v) formic acid in acetonitrile (B), with a gradient of 0.1–9% (B) at 0.25–0.4 min, 9–17% (B) at 0.4–0.8 min, 17–99.9% (B) at 0.8–1.9 min, 99.9–0.1% (B) at 1.9–2.1 min, and 0.1% (B) at 2.11–2.7 min. The flow rate was 0.24 mL/min, and the injection volume was 1 $\mu$L.

The mass spectrometer was operated in both positive and negative ESI modes. Key parameters included: nebulizing gas flow 3 mL/min, heating gas 10 L/min, drying gas 10 L/min, interface temperature 300 °C, dilution line 250 °C, heat block 400 °C, CID gas 270 kPa, and capillary voltages of +4 kV and –3.5 kV. Data were acquired in scheduled multiple reaction monitoring (MRM) mode for MS/MS detection, and compound-specific transitions are provided in Supplementary Table 1.

Raw data (lcd files) were converted to Abf format using the Reifycs Abf Converter (https://www.reifycs.com/AbfConverter/). Peak areas of LC–QqQ–MS data were quantified using MRMPROBS software (http://prime.psc.riken.jp/compms/mrmprobs/main.html), and peak areas were normalized based on the internal standards.

#### A.4 Microbiome Analysis

##### DNA extraction

Genomic DNA was isolated from root and rhizosphere soil samples following previously reported procedures. 100 mg of finely powdered sample was transferred into a 1.5 mL microtube and flash-frozen in liquid nitrogen. Subsequently, 200 µL of lysis/binding buffer (LBB) containing 1 M LiCl (Cat. #L7026-500ML, Sigma-Aldrich, St. Louis, USA), 100 mM Tris-HCl (Cat. #318-90225, Wako Pure Chemical Corporation, Osaka, Japan), 1% SDS (Cat. #313-90275, Wako Pure Chemical Corporation), 10 mM EDTA, pH 8.0 (Cat. #311-90075, Wako Pure Chemical Corporation), Antifoam A (Cat. #A5633-25G, Sigma-Aldrich), 5 mM dithiothreitol (Cat. #048-29224, Wako Pure Chemical Corporation), 11.2 M 3-mercapto-1,2-propanediol (Cat. #139-16452, Wako Pure Chemical Corporation), and DNase/RNase-free water (Cat. #10977015, Thermo Fisher Scientific, Waltham, MA, USA) was added. The mixture was vortexed briefly and incubated for 5 min at room temperature (24 °C).

Samples were centrifuged at 13,000 rpm for 10 min at room temperature, and 50 µL of the supernatant was combined with an equal volume of AMPure XP beads (Cat. #A63881, Beckman Coulter, Inc.) for DNA purification and size selection. The mixture was vortexed, incubated at room temperature for 5 min, and placed on a magnetic rack for another 5 min. After discarding the supernatant, beads were washed twice with 200 µL of 80% ethanol. DNA was eluted from the beads using 15 µL of 10 mM Tris-HCl (pH 7.5).

##### Bacterial 16S rRNA gene sequencing

Amplicon libraries were generated using a modified two-step PCR amplification protocol. For the initial PCR, the V4 region of the bacterial 16S rRNA gene was amplified using primers (515f: 5'- TCG TCG GCA GCG TCA GAT GTG TAT AAG AGA

CAG – [3–6-mer Ns] – GTG YCA GCM GCC GCG GTA A -3'; 806rB: 5'- GTC TCG TGG GCT CGG AGA TGT GTA TAA GAG ACA G – [3–6-mer Ns] – GGA CTA CNV GGG TWT CTA AT -3'). PCR reactions (10 µL) contained 1 µL of ten-fold diluted DNA template, 2× KAPA HiFi HotStart ReadyMix (Cat. #07958935001, KAPA Biosystems Inc., USA), 0.2 µM of each primer, and 1 µM blocking primers (mPNA and pPNA; PNA BIO, Inc., Newbury Park, CA, USA). Thermal cycling conditions were: initial denaturation at 95 °C for 3 min; 35 cycles of 98 °C for 20 s, 78 °C for 10 s, 50 °C for 30 s, and 72 °C for 90 s; followed by a final extension at 72 °C for 5 min (ramp rate = 1 °C/s).

PCR products were purified using ExoSAP-IT Express (Cat. #75001.1.EA; Thermo Fisher Scientific). Five microliters of the PCR product were mixed with 2 µL of ExoSAP-IT Express, incubated at 37 °C for 4 min, and heat-inactivated at 80 °C for 1 min.

The second PCR incorporated Illumina index sequences using primers: forward (5'- AAT GAT ACG GCG ACC ACC GAG ATC TAC AC – [8-mer index] – TCG TCG GCA GCG TC -3') and reverse (5'- CAA GCA GAA GAC GGC ATA CGA GAT – [8-mer index] – GTC TCG TGG GCT CGG -3'). Reactions (10 µL) included 0.8 µL of purified first-round amplicons, 2× KAPA HiFi HotStart ReadyMix, 1 µM of each index primer, and 1 µM blocking primers. Cycling conditions were: 95 °C for 3 min; 8 cycles of 98 °C for 20 s, 78 °C for 10 s, 55 °C for 30 s, and 72 °C for 90 s; and a final extension at 72 °C for 5 min (ramp rate = 1 °C/s).

Amplicons were purified using AMPure XP beads, incubated for 5 min at room temperature, and placed on a magnetic rack. After two washes with 200 µL of 80% ethanol, DNA was eluted with 15 µL of 10 mM Tris-HCl (pH 7.5). Library concentrations were measured using the Quant-iT PicoGreen dsDNA assay (Cat. #P7581, Invitrogen) on a microplate reader (Infinite 200 PRO M Nano+, TECAN Japan Co., Ltd., Kanagawa, Japan). Libraries were normalized and pooled in equimolar amounts, quantified by qPCR using the NEBNext Library Quant Kit for Illumina (Cat. #E7630L, New England BioLabs, USA), and diluted to 3 nM. The denatured library pool was supplemented with 20% PhiX Control v3 (Cat. #FC-110-3001, Illumina, USA) and sequenced on an Illumina MiSeq platform with the MiSeq Reagent Kit v3 (2×300 bp, Illumina).

#### 16S V4 rRNA data preprocessing

Raw paired-end reads were analyzed with QIIME2 (ver. 2020.6.0). FASTQ files were imported into QIIME2, and adapter sequences were removed with the Cutadapt plugin. For ASV inference, reads were processed with the DADA2 pipeline, which included truncation (forward reads at 240 bp, reverse reads at 180 bp), quality filtering, denoising, merging of read pairs, and chimera removal. Taxonomic assignment of amplicon sequence variants (ASVs) was performed using the SILVA database (ver. 138). Sequences assigned to Archaea, Eukaryota, mitochondria, or chloroplasts were excluded from downstream analyses.

##### A.5 Multi-Omics Integration and Scaling

For Random Forest (RF) modeling, the genome, metabolome, and microbiome features were concatenated into a single input matrix without enforcing equal feature numbers across omics layers. This integration strategy was adopted to support the primary objective of this study, which was to investigate the coordination and interaction patterns among different omics layers, rather than to compare or optimize predictive performance.

As the number of genome-wide SNP markers was substantially larger than that of the other omics layers, linkage disequilibrium (LD) pruning was applied as a standard genome analysis procedure. This reduced the original set of 425,858 SNPs to 3,078 representative markers (LD threshold = 0.001), which were broadly distributed across the genome, providing approximate genome-wide coverage and a certain prediction performance (Figs. B1,B2).

Microbiome features were scaled prior to modeling, and variables with missing values or zero variance within each condition were removed by filtering. The control and drought-treated samples were preprocessed using an identical pipeline to ensure consistency. Metabolome features were similarly scaled, and all variables that passed the quality control were retained.

Differences in feature dimensionality and scale across omics layers were not explicitly corrected. Enforcing an artificial balance among omics types would require imposing arbitrary constraints and restricting the diversity of the biological signals captured. Instead, RF was used to allow for intrinsic data-driven feature selection based on the feature importance scores.

Informative features from all omics layers were retained in the downstream analyses, indicating that no single omics layer dominated the model solely because of feature dimensionality. The comparison with BLUP supported the fact that RF exhibited stable behavior across heterogeneous omics scales (Fig. 1), which is consistent with the study's emphasis on cross-omics coordination and interaction analyses.

### Appendix B Supplementary Figures and Table

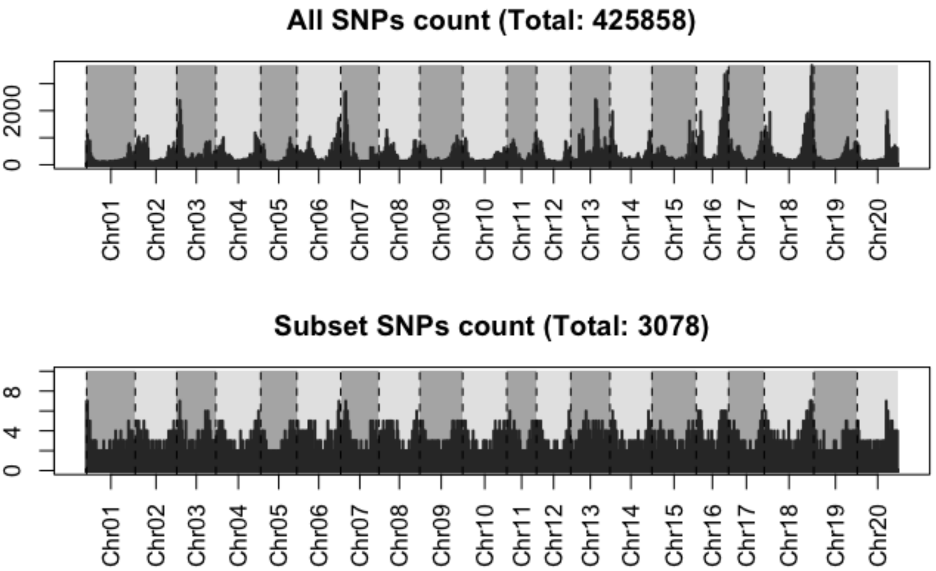


**Fig. B1**: SNPs coverage comparison. The SNP mapping in the full set of 425,838 SNPs ( LD threshold of 0.95) and a subset of 3,078 SNPs filtered by a LD threshold of 0.001.

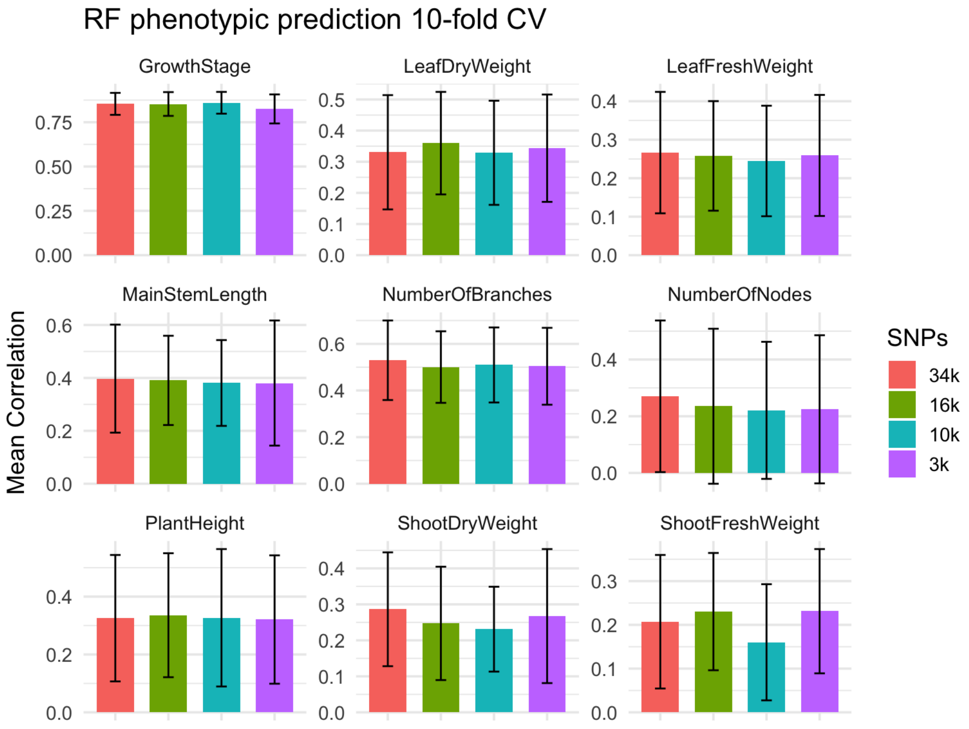


**Fig. B2**: Mean Pearson correlation (±1 SD) from 10-fold cross-validation of RF models predicting phenotypes. Bars compare SNP marker sets of different sizes; 34k (34,632), 16k(16,419), 10k(10,143), and 3k(3,078) loci, respectively. The legend shows the number of markers.

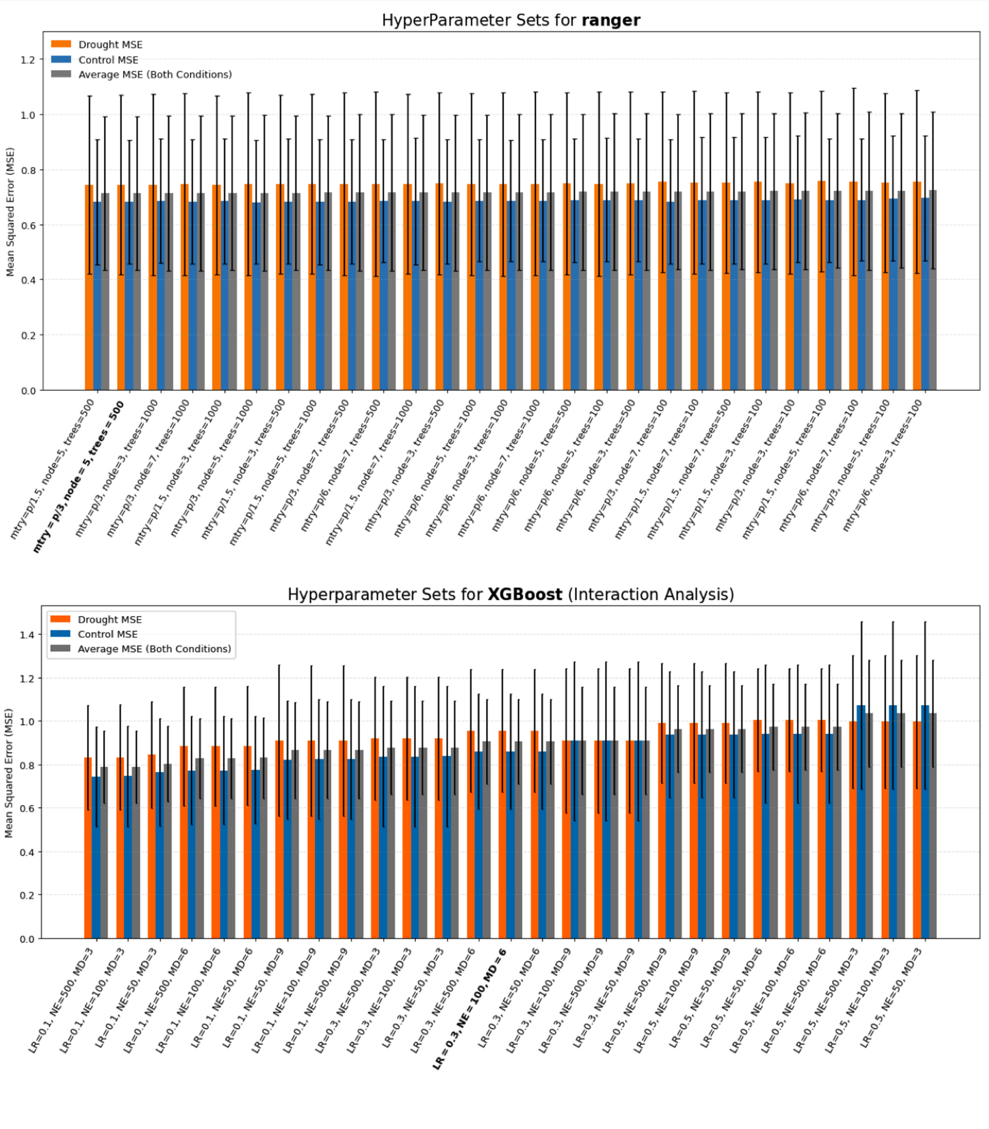


**Fig. B3**: Hyperparameter tuning for phenotypic prediction under drought and control conditions. Top: Ranger, Bottom: XGBoost. Each bar represents the mean squared error (MSE) averaged over 10-fold cross-validation, with error bars indicating the standard deviation across folds for a specific combination of three hyperparameters. Orange bars represent MSE under drought conditions, blue barsrepresent MSE under control conditions, and gray bars show the average MSE across both conditions. The hyperparameter combinations selected for this study are highlighted in bold.

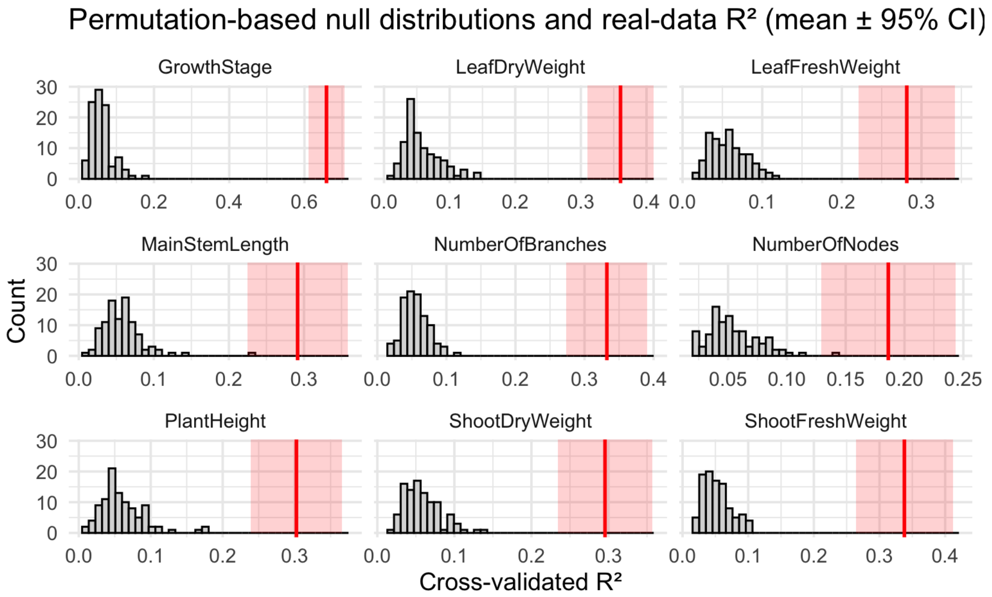


**Fig. B4**: Null distribution of the cross-validated coefficient of determination ($R^2$) obtained by permuting the phenotypic labels (100 permutations, 10-fold cross-validation each) in phenotypic prediction by RF model (input: genome, microbiome, and metabolome). The gray histogram represents the null distribution under the assumption of no association between predictors and phenotype. The vertical red line indicates the mean $R^2$ obtained from repeated 5-fold cross-validation using the real data, and the shaded region denotes its 95% confidence interval (mean $\pm$ 1.96 SE). The real-data $R^2$ lies far outside the null distribution, demonstrating that the predictive performance cannot be explained by random associations.

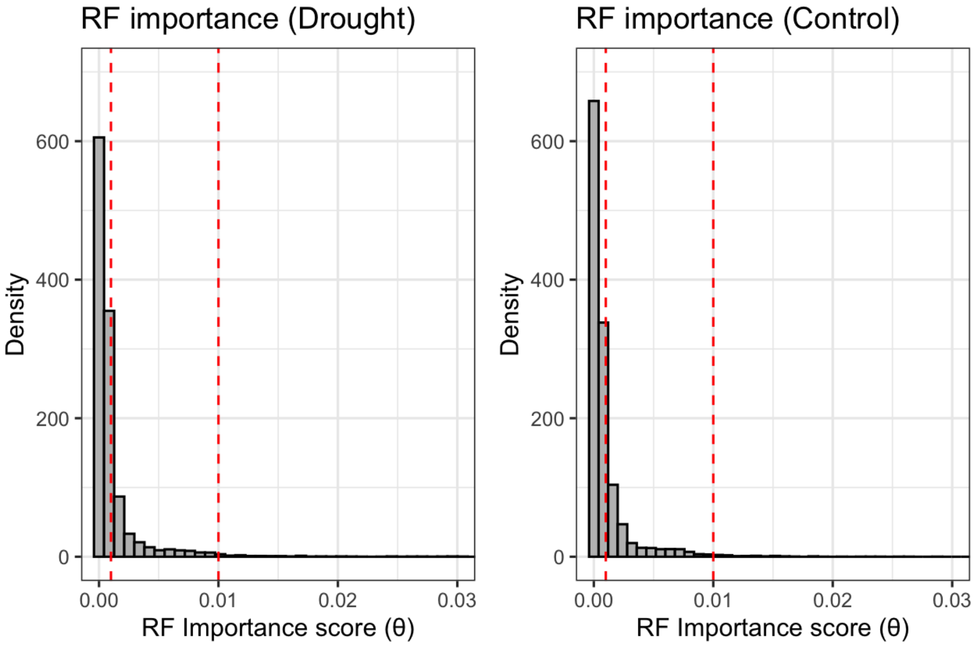


**Fig. B5**: Distribution of RF importance value $\theta$. The vertical lines show the threshold ($\theta$=0.001,0.01). Approximately 1% and 25% of features are retained at these thresholds, respectively.

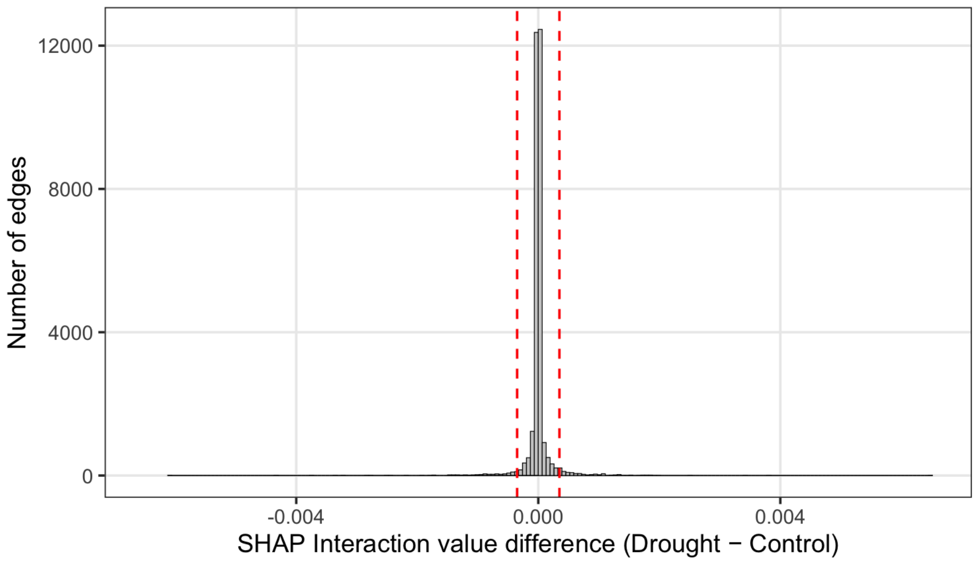


**Fig. B6**: Distribution of SHAP interaction value differences (Drought − Control) across all feature pairs. Only non-zero interaction edges are shown (5,563 of 3,716,793). Red dashed vertical lines indicate the thresholds corresponding to $Z = \pm 30$, derived from the pooled null distribution.

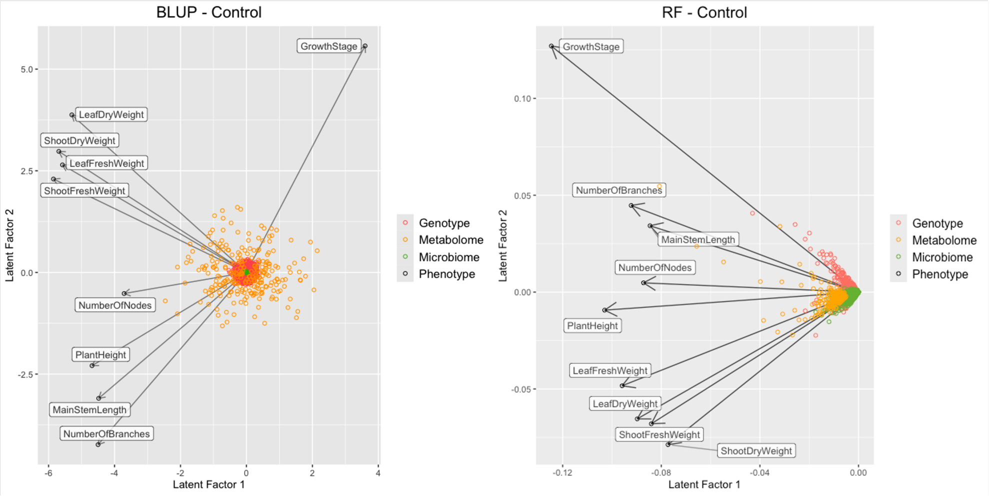


**Fig. B7**: Feature-trait structures derived from BLUP and RF in predicting biomass-related phenotypes under control conditions. Feature importance matrices from both models were subjected to singular value decomposition (SVD), and the first two latent factors are visualized. Points represent features and phenotypes projected into the latent space. Color indicates data type: genotype, metabolome, or microbiome. Vectors for phenotypes are shown as arrows.

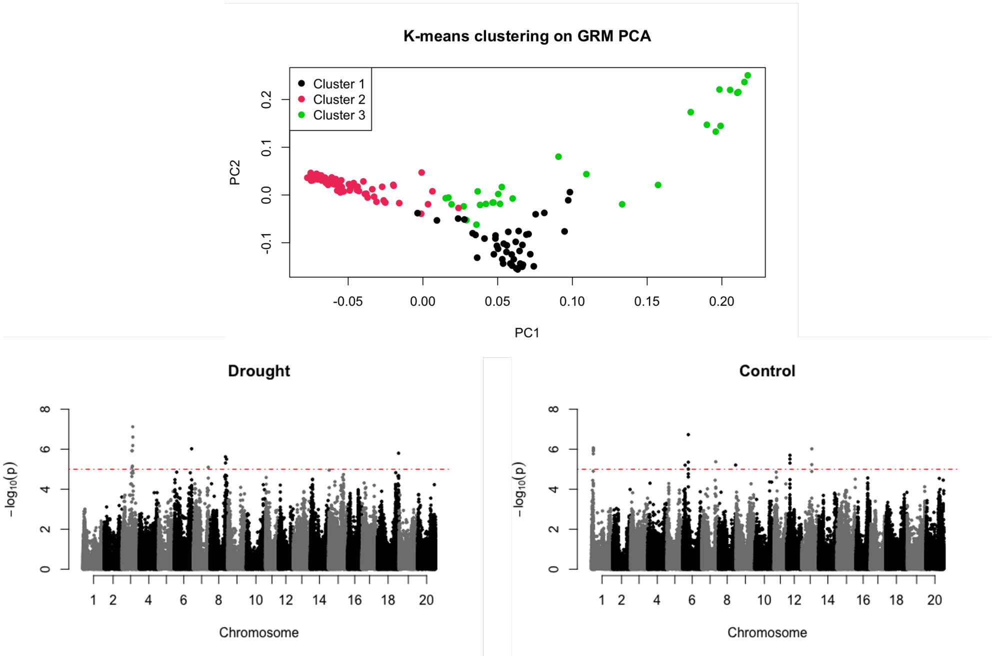


**Fig. B8**: The genomic clustering and GWAS results. Top plot: PCA (principal component analysis) and k-means clustering of genomic relationship matrix (covariance matrix of SNP markers). Bottom plot: Results of GWAS. The red horizontal dotted line indicates the threshold of $p < 1 \times 10^{-5}$. GWAS was performed by incorporating the three principal components.

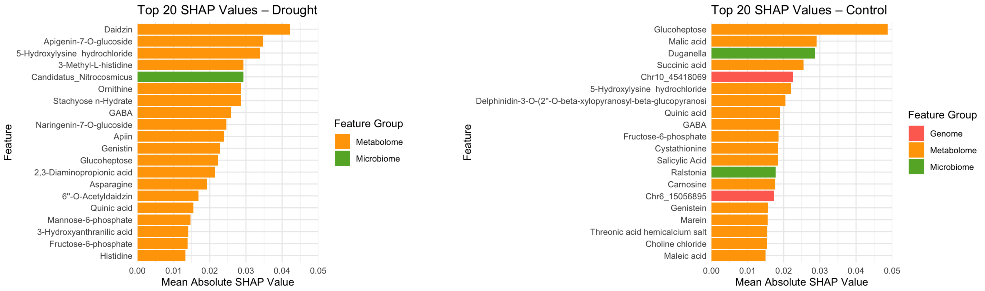


**Fig. B9**: Top 20 SHAP values in drought and control conditions.

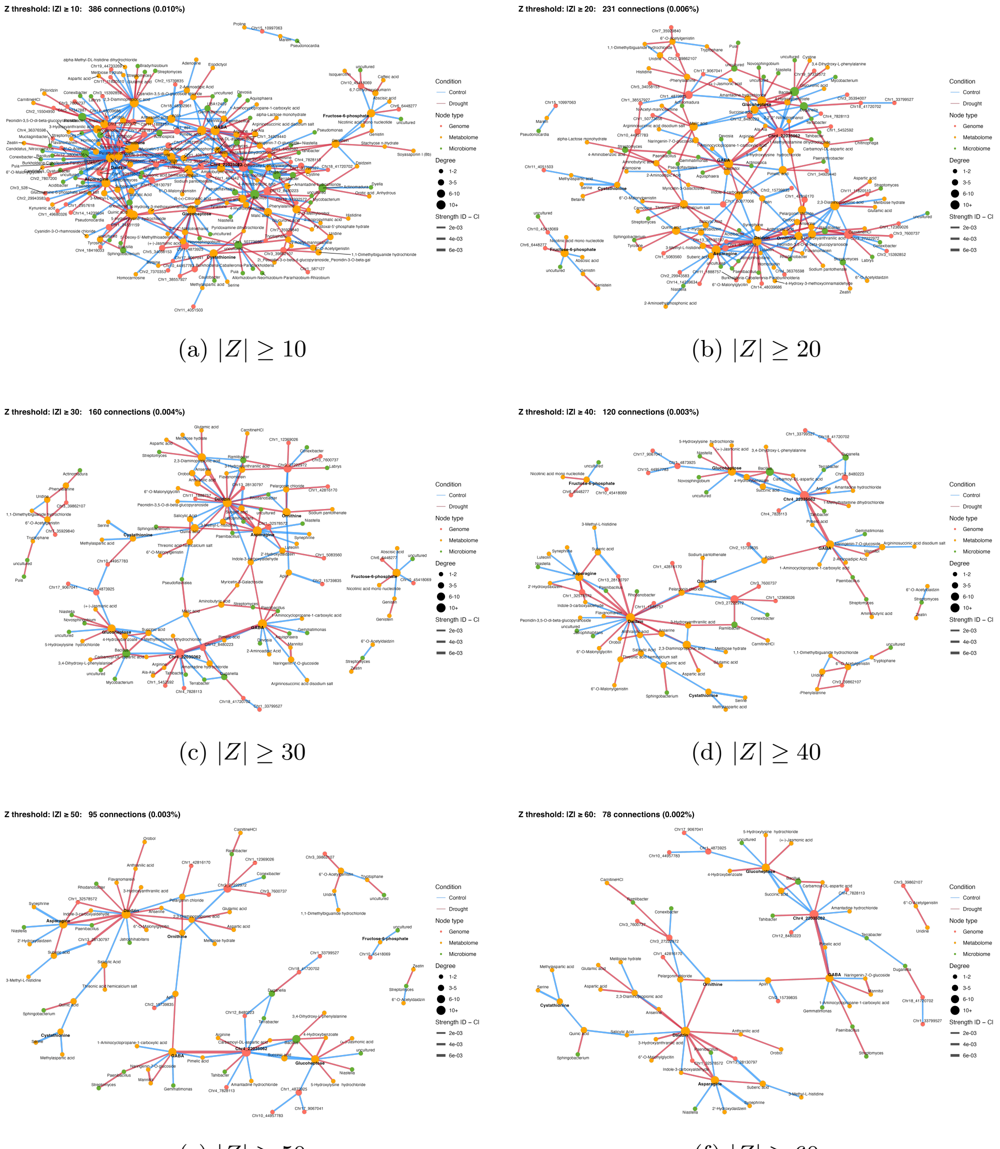


(a) $|Z| \geq 10$ (b) $|Z| \geq 20$

(c) $|Z| \geq 30$ (d) $|Z| \geq 40$

(e) $|Z| \geq 50$ (f) $|Z| \geq 60$

**Fig. B10**: Condition-specific SHAP interaction difference networks under varying Z-score thresholds. Networks were constructed from the differential SHAP interaction matrix (drought minus control), retaining only edges with large absolute Z-scores. Increasing the Z threshold progressively restricts the network to more extreme, environment-specific interactions.

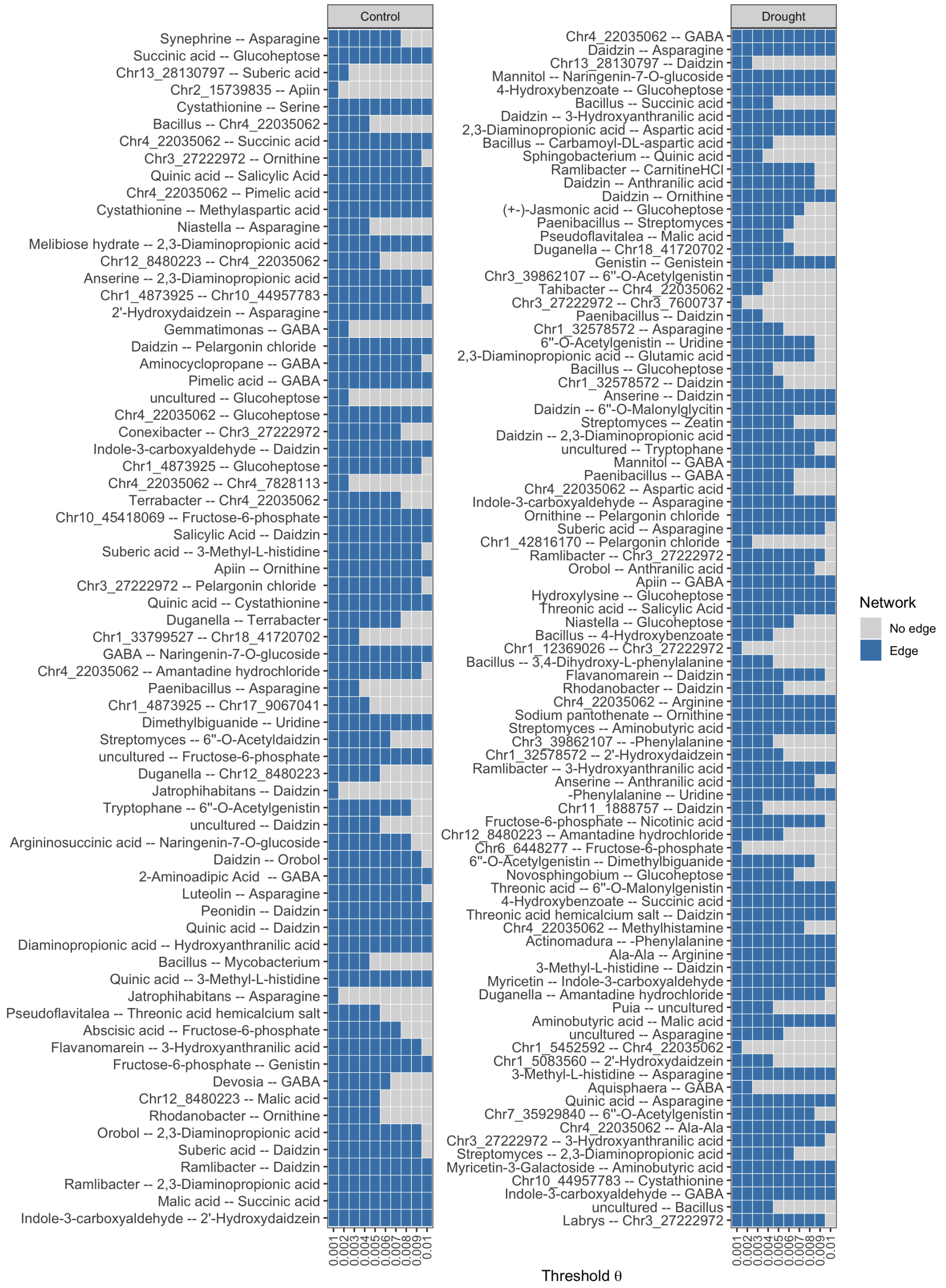


**Fig. B11**: Sensitivity analysis of the SHAP interaction network to the Random Forest importance threshold $\theta$. The SHAP interaction network was first constructed under the most permissive threshold ($\theta = 0.001$), and the retained edge set was then re-evaluated by restricting the network to nodes that remained selected under increasingly stringent $\theta$ values. For each threshold, an edge was retained only when both of its incident nodes were still included after RF-based feature filtering. The two panels show drought-specific and control-specific subnetworks, respectively, and each row corresponds to one previously identified edge with $|Z| \geq 30$.

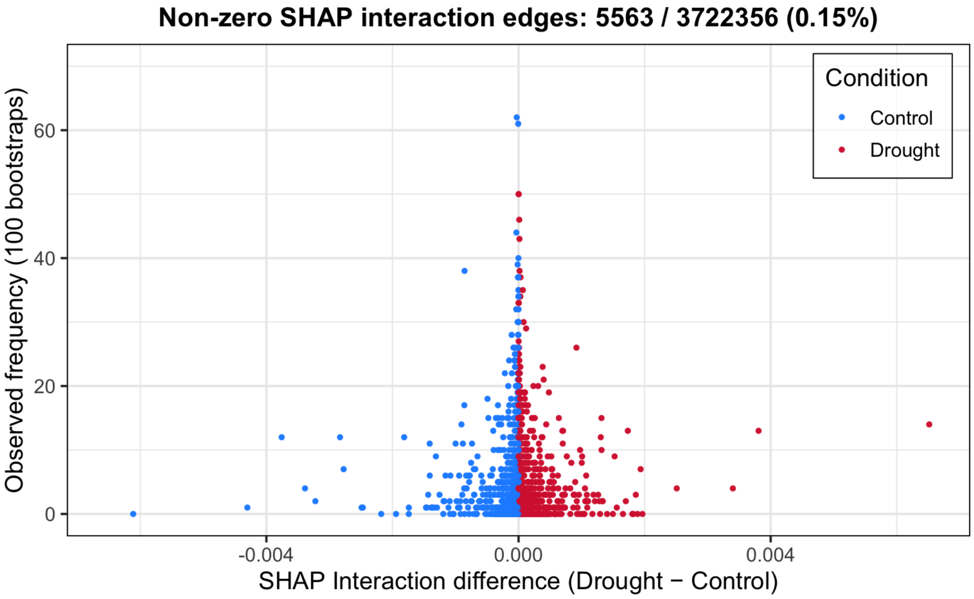


**Fig. B12**: Stability of SHAP interaction differences between drought and control. Each point represents one feature–feature interaction edge. The x-axis shows the signed difference in SHAP interaction strength (Drought − Control) from the point estimate, and the y-axis shows the number of bootstrap replicates (out of 100) in which the edge was observed as non-zero (5563/3722356; 0.15 %). Colors indicate the direction of the point-estimate difference (positive: drought-enriched; negative: control-enriched).

| $\theta$ | Number of Features | Number of Edges | (Ratio) |
|---|---|---|---|
| 0.001 | 2729 | 160 | (1.00) |
| 0.002 | 1341 | 153 | (0.96) |
| 0.003 | 877 | 146 | (0.91) |
| 0.004 | 677 | 140 | (0.88) |
| 0.005 | 539 | 126 | (0.79) |
| 0.006 | 454 | 113 | (0.71) |
| 0.007 | 361 | 103 | (0.64) |
| 0.008 | 277 | 96 | (0.60) |
| 0.009 | 216 | 85 | (0.53) |
| 0.010 | 159 | 66 | (0.41) |

**Table B1**: Effect of the threshold $\theta$ on the number of selected features and remaining edges. Here, $\theta$ denotes the threshold applied to the RF importance scores. The column "Number of Features" represents the number of features whose importance exceeds the threshold. The SHAP interaction network was then restricted to these selected features, and "Number of Edges" denotes the number of edges in the resulting SHAP subnetwork ($|Z| \geq 30$). The ratio represents the proportion of remaining edges relative to the baseline network obtained at $\theta = 0.001$ (160 edges).

| target_label | partner_label | weight | z | p-value | Stability |
|---|---|---|---|---|---|
| Chr4_22035062 | GABA | 6.5e-03 | 559.45 | 9.9e-03 | 14 |
| Asparagine | Daidzin | 3.8e-03 | 326.99 | 9.9e-03 | 13 |
| Chr13_28130797 | Daidzin | 3.4e-03 | 291.96 | 9.9e-03 | 4 |
| Naringenin-7-O-glucoside | Mannitol | 2.5e-03 | 215.48 | 9.9e-03 | 4 |
| Glucoheptose | 4-Hydroxybenzoate | 1.9e-03 | 166.31 | 9.9e-03 | 7 |
| Bacillus | Succinic acid | 1.9e-03 | 159.76 | 9.9e-03 | 3 |
| 3-Hydroxyanthranilic acid | Daidzin | 1.7e-03 | 148.86 | 9.9e-03 | 13 |
| 2,3-Diaminopropionic acid | Aspartic acid | 1.7e-03 | 146.13 | 9.9e-03 | 2 |
| Bacillus | Carbamoyl-DL-aspartic acid | 1.5e-03 | 130.88 | 2.0e-02 | 9 |
| Sphingobacterium | Quinic acid | 1.3e-03 | 114.72 | 9.9e-03 | 2 |
| Ramlibacter | CarnitineHCl | 1.3e-03 | 114.12 | 9.9e-03 | 2 |
| Anthranilic acid | Daidzin | 1.3e-03 | 113.07 | 9.9e-03 | 10 |
| Ornithine | Daidzin | 1.3e-03 | 112.99 | 2.0e-02 | 15 |
| Glucoheptose | (+-)-Jasmonic acid | 1.3e-03 | 112.00 | 9.9e-03 | 12 |
| Streptomyces | Paenibacillus | 1.3e-03 | 110.09 | 9.9e-03 | 2 |
| Pseudoflavitalea | Malic acid | 1.3e-03 | 108.44 | 9.9e-03 | 3 |
| Duganella | Chr18_41720702 | 1.2e-03 | 104.01 | 9.9e-03 | 2 |
| Genistin | Genistein | 1.2e-03 | 102.08 | 9.9e-03 | 3 |
| 6”-O-Acetylgenistin | Chr3_39862107 | 1.1e-03 | 94.72 | 9.9e-03 | 2 |
| Chr4_22035062 | Tahibacter | 1.1e-03 | 93.01 | 9.9e-03 | 3 |
| Chr3_7600737 | Chr3_27222972 | 1.1e-03 | 92.17 | 9.9e-03 | 2 |
| Paenibacillus | Daidzin | 1.1e-03 | 92.10 | 9.9e-03 | 6 |
| Chr1_32578572 | Asparagine | 1.1e-03 | 91.76 | 9.9e-03 | 2 |
| Uridine | 6”-O-Acetylgenistin | 1.1e-03 | 90.91 | 9.9e-03 | 2 |
| 2,3-Diaminopropionic acid | Glutamic acid | 1.1e-03 | 90.53 | 9.9e-03 | 4 |
| Bacillus | Glucoheptose | 1.0e-03 | 86.63 | 9.9e-03 | 9 |
| Chr1_32578572 | Daidzin | 1.0e-03 | 86.12 | 9.9e-03 | 8 |
| Anserine | Daidzin | 9.8e-04 | 83.84 | 9.9e-03 | 10 |
| Daidzin | 6”-O-Malonylglycitin | 9.6e-04 | 82.24 | 9.9e-03 | 3 |
| Streptomyces | Zeatin | 9.2e-04 | 79.34 | 9.9e-03 | 2 |
| 2,3-Diaminopropionic acid | Daidzin | 9.2e-04 | 78.85 | 9.9e-03 | 26 |
| uncultured | Tryptophane | 9.0e-04 | 77.19 | 9.9e-03 | 5 |
| GABA | Mannitol | 8.9e-04 | 76.78 | 9.9e-03 | 3 |
| Paenibacillus | GABA | 8.8e-04 | 75.35 | 9.9e-03 | 3 |
| Chr4_22035062 | Carbamoyl-DL-aspartic acid | 8.8e-04 | 75.27 | 9.9e-03 | 3 |
| Asparagine | Indole-3-carboxyaldehyde | 8.4e-04 | 71.89 | 9.9e-03 | 8 |
| Ornithine | Pelargonin chloride | 8.2e-04 | 70.49 | 9.9e-03 | 4 |
| Asparagine | Suberic acid | 7.7e-04 | 65.74 | 9.9e-03 | 9 |
| Chr1_42816170 | Pelargonin chloride | 7.4e-04 | 63.95 | 9.9e-03 | 4 |
| Chr3_27222972 | Ramlibacter | 7.4e-04 | 63.13 | 9.9e-03 | 3 |
| Anthranilic acid | Orobol | 7.3e-04 | 62.65 | 9.9e-03 | 3 |
| GABA | Apiin | 7.1e-04 | 60.63 | 9.9e-03 | 13 |
| Glucoheptose | 5-Hydroxylysine hydrochloride | 7.0e-04 | 60.25 | 9.9e-03 | 11 |
| Threonic acid hemicalcium salt | Salicylic Acid | 7.0e-04 | 59.70 | 9.9e-03 | 13 |
| Niastella | Glucoheptose | 6.9e-04 | 59.53 | 9.9e-03 | 3 |
| Bacillus | 4-Hydroxybenzoate | 6.9e-04 | 59.34 | 9.9e-03 | 2 |
| Chr1_12369026 | Chr3_27222972 | 6.4e-04 | 54.89 | 9.9e-03 | 15 |
| Bacillus | 3,4-Dihydroxy-L-phenylalanine | 6.4e-04 | 54.76 | 9.9e-03 | 2 |
| Flavanomarein | Daidzin | 6.3e-04 | 54.10 | 9.9e-03 | 4 |
| Rhodanobacter | Daidzin | 6.2e-04 | 53.57 | 9.9e-03 | 6 |
| Chr4_22035062 | Arginine | 6.2e-04 | 52.98 | 9.9e-03 | 2 |
| Ornithine | Sodium pantothenate | 5.6e-04 | 48.08 | 9.9e-03 | 5 |
| Streptomyces | Aminobutyric acid | 5.6e-04 | 47.96 | 9.9e-03 | 4 |
| -Phenylalanine | Chr3_39862107 | 5.5e-04 | 46.81 | 9.9e-03 | 4 |
| Chr1_32578572 | 2’-Hydroxydaidzein | 5.4e-04 | 46.75 | 9.9e-03 | 3 |
| Ramlibacter | 3-Hydroxyanthranilic acid | 5.4e-04 | 46.48 | 9.9e-03 | 4 |
| Anthranilic acid | Anserine | 5.4e-04 | 46.44 | 9.9e-03 | 6 |
| -Phenylalanine | Uridine | 5.3e-04 | 45.93 | 9.9e-03 | 7 |
| Chr11_1888757 | Daidzin | 5.2e-04 | 44.79 | 9.9e-03 | 7 |
| Nicotinic acid mono nucleotide | Fructose-6-phosphate | 5.2e-04 | 44.68 | 9.9e-03 | 9 |
| Amantadine hydrochloride | Chr12_8480223 | 5.1e-04 | 43.96 | 9.9e-03 | 5 |
| Chr6_6448277 | Fructose-6-phosphate | 5.1e-04 | 43.55 | 9.9e-03 | 3 |
| 1,1-Dimethylbiguanide hydrochloride | 6”-O-Acetylgenistin | 5.1e-04 | 43.54 | 9.9e-03 | 5 |
| Novosphingobium | Glucoheptose | 4.9e-04 | 42.07 | 9.9e-03 | 2 |
| Threonic acid hemicalcium salt | 6”-O-Malonylgenistin | 4.9e-04 | 42.07 | 9.9e-03 | 5 |
| Succinic acid | 4-Hydroxybenzoate | 4.8e-04 | 41.64 | 9.9e-03 | 3 |
| Threonic acid hemicalcium salt | Daidzin | 4.8e-04 | 41.35 | 9.9e-03 | 19 |
| Chr4_22035062 | 1-Methylhistamine dihydrochloride | 4.7e-04 | 40.23 | 9.9e-03 | 4 |
| Actinomadura | -Phenylalanine | 4.6e-04 | 39.73 | 9.9e-03 | 9 |
| Ala-Ala | Arginine | 4.4e-04 | 37.70 | 9.9e-03 | 3 |
| 3-Methyl-L-histidine | Daidzin | 4.4e-04 | 37.69 | 9.9e-03 | 14 |
| Myricetin-3-Galactoside | Indole-3-carboxyaldehyde | 4.3e-04 | 36.73 | 9.9e-03 | 3 |
| Duganella | Amantadine hydrochloride | 4.3e-04 | 36.59 | 9.9e-03 | 7 |
| uncultured | Puia | 4.2e-04 | 36.06 | 9.9e-03 | 3 |
| Aminobutyric acid | Malic acid | 4.1e-04 | 34.90 | 9.9e-03 | 3 |
| uncultured | Asparagine | 4.0e-04 | 34.77 | 9.9e-03 | 3 |
| Chr1_5452592 | Chr4_22035062 | 4.0e-04 | 34.20 | 9.9e-03 | 21 |
| Chr1_5083560 | 2’-Hydroxydaidzein | 3.8e-04 | 32.83 | 9.9e-03 | 23 |
| Asparagine | 3-Methyl-L-histidine | 3.8e-04 | 32.55 | 9.9e-03 | 12 |
| Aquisphaera | GABA | 3.7e-04 | 32.02 | 9.9e-03 | 2 |
| Asparagine | Quinic acid | 3.7e-04 | 32.00 | 2.0e-02 | 2 |
| 6”-O-Acetylgenistin | Chr7_35929840 | 3.7e-04 | 31.97 | 9.9e-03 | 2 |
| Chr4_22035062 | Ala-Ala | 3.7e-04 | 31.89 | 9.9e-03 | 5 |
| Chr3_27222972 | 3-Hydroxyanthranilic acid | 3.7e-04 | 31.46 | 9.9e-03 | 12 |
| Streptomyces | 2,3-Diaminopropionic acid | 3.6e-04 | 31.33 | 9.9e-03 | 2 |
| Myricetin-3-Galactoside | Aminobutyric acid | 3.6e-04 | 30.97 | 9.9e-03 | 4 |
| Chr10_44957783 | Cystathionine | 3.6e-04 | 30.94 | 9.9e-03 | 2 |
| GABA | Indole-3-carboxyaldehyde | 3.6e-04 | 30.72 | 2.0e-02 | 8 |
| uncultured | Bacillus | 3.6e-04 | 30.49 | 9.9e-03 | 5 |
| Chr3_27222972 | Labrys | 3.5e-04 | 30.05 | 9.9e-03 | 7 |

**Table B2**: SHAP interaction edges with positive Z-scores ($Z \geq 30$). *weight* denotes the interaction difference $|\Delta|$ (Drought − Control), with two-sided $p$-values and bootstrap stability.

| target_label | partner_label | weight | z | p-value | Stability |
|---|---|---|---|---|---|
| Asparagine | Synephrine | 3.8e-03 | -322.72 | 9.9e-03 | 12 |
| Succinic acid | Glucoheptose | 3.4e-03 | -290.88 | 2.0e-02 | 4 |
| Chr13_28130797 | Suberic acid | 3.2e-03 | -276.77 | 9.9e-03 | 2 |
| Chr2_15739835 | Apiin | 2.8e-03 | -243.16 | 9.9e-03 | 12 |
| Cystathionine | Serine | 2.8e-03 | -238.25 | 9.9e-03 | 7 |
| Chr4_22035062 | Bacillus | 1.8e-03 | -155.80 | 9.9e-03 | 12 |
| Chr4_22035062 | Succinic acid | 1.4e-03 | -123.08 | 9.9e-03 | 3 |
| Chr3_27222972 | Ornithine | 1.4e-03 | -121.02 | 9.9e-03 | 11 |
| Quinic acid | Salicylic Acid | 1.4e-03 | -120.89 | 9.9e-03 | 6 |
| Chr4_22035062 | Pimelic acid | 1.3e-03 | -112.57 | 9.9e-03 | 9 |
| Cystathionine | Methylaspartic acid | 1.3e-03 | -107.60 | 9.9e-03 | 3 |
| Niastella | Asparagine | 1.2e-03 | -102.17 | 9.9e-03 | 2 |
| 2,3-Diaminopropionic acid | Melibiose hydrate | 1.2e-03 | -100.95 | 9.9e-03 | 2 |
| Chr4_22035062 | Chr12_8480223 | 1.2e-03 | -99.56 | 9.9e-03 | 6 |
| 2,3-Diaminopropionic acid | Anserine | 1.1e-03 | -93.72 | 9.9e-03 | 2 |
| Chr1_4873925 | Chr10_44957783 | 1.1e-03 | -92.84 | 9.9e-03 | 6 |
| Asparagine | 2'-Hydroxydaidzein | 1.0e-03 | -85.78 | 9.9e-03 | 11 |
| Gemmatimonas | GABA | 9.8e-04 | -84.33 | 9.9e-03 | 2 |
| Pelargonin chloride | Daidzin | 9.4e-04 | -80.95 | 9.9e-03 | 6 |
| 1-Aminocyclopropane-1-carboxylic acid | GABA | 9.2e-04 | -79.34 | 9.9e-03 | 2 |
| GABA | Pimelic acid | 9.1e-04 | -77.77 | 9.9e-03 | 14 |
| uncultured | Glucoheptose | 8.9e-04 | -76.64 | 9.9e-03 | 3 |
| Chr4_22035062 | Glucoheptose | 8.8e-04 | -75.83 | 9.9e-03 | 11 |
| Chr3_27222972 | Conexibacter | 8.6e-04 | -74.12 | 9.9e-03 | 4 |
| Indole-3-carboxyaldehyde | Daidzin | 8.6e-04 | -73.86 | 9.9e-03 | 17 |
| Chr1_4873925 | Glucoheptose | 8.6e-04 | -73.55 | 9.9e-03 | 38 |
| Chr4_7828113 | Chr4_22035062 | 8.5e-04 | -73.08 | 9.9e-03 | 4 |
| Chr4_22035062 | Terrabacter | 8.4e-04 | -71.76 | 9.9e-03 | 2 |
| Chr10_45418069 | Fructose-6-phosphate | 8.3e-04 | -71.41 | 9.9e-03 | 6 |
| Salicylic Acid | Daidzin | 8.2e-04 | -70.70 | 2.0e-02 | 4 |
| Suberic acid | 3-Methyl-L-histidine | 7.9e-04 | -68.07 | 9.9e-03 | 6 |
| Ornithine | Apiin | 7.8e-04 | -67.30 | 9.9e-03 | 5 |
| Chr3_27222972 | Pelargonin chloride | 7.7e-04 | -66.38 | 9.9e-03 | 6 |
| Cystathionine | Quinic acid | 7.5e-04 | -64.61 | 9.9e-03 | 8 |
| Duganella | Terrabacter | 7.5e-04 | -64.56 | 9.9e-03 | 2 |
| Chr1_33799527 | Chr18_41720702 | 7.4e-04 | -63.74 | 9.9e-03 | 2 |
| GABA | Naringenin-7-O-glucoside | 7.4e-04 | -63.35 | 9.9e-03 | 11 |
| Chr4_22035062 | Amantadine hydrochloride | 7.1e-04 | -60.96 | 9.9e-03 | 7 |
| Paenibacillus | Asparagine | 7.0e-04 | -60.52 | 9.9e-03 | 3 |
| Chr1_4873925 | Chr17_9067041 | 7.0e-04 | -60.24 | 9.9e-03 | 3 |
| Chr1_10557956 | Guanosine | 6.9e-04 | -58.88 | 9.9e-03 | 7 |
| 1,1-Dimethylbiguanide hydrochloride | Uridine | 6.5e-04 | -55.94 | 9.9e-03 | 9 |
| Streptomyces | 6"-O-Acetyldaidzin | 6.3e-04 | -53.75 | 9.9e-03 | 2 |
| uncultured | Fructose-6-phosphate | 6.1e-04 | -52.51 | 9.9e-03 | 2 |
| Duganella | Chr12_8480223 | 6.1e-04 | -52.14 | 9.9e-03 | 2 |
| Jatrophihabitans | Daidzin | 5.9e-04 | -50.75 | 9.9e-03 | 6 |
| Tryptophane | 6"-O-Acetylgenistin | 5.9e-04 | -50.28 | 9.9e-03 | 4 |
| uncultured | Daidzin | 5.7e-04 | -48.83 | 9.9e-03 | 5 |
| Argininosuccinic acid disodium salt | Naringenin-7-O-glucoside | 5.6e-04 | -48.43 | 9.9e-03 | 3 |
| Daidzin | Orobol | 5.2e-04 | -44.65 | 9.9e-03 | 2 |
| GABA | 2-Aminoadipic Acid | 5.1e-04 | -43.87 | 9.9e-03 | 4 |
| Asparagine | Luteolin | 5.1e-04 | -43.76 | 9.9e-03 | 2 |
| Peonidin-3,5-O-di-beta-glucopyranoside | Daidzin | 5.1e-04 | -43.54 | 9.9e-03 | 3 |
| Quinic acid | Daidzin | 4.9e-04 | -42.25 | 9.9e-03 | 18 |
| 2,3-Diaminopropionic acid | 3-Hydroxyanthranilic acid | 4.8e-04 | -41.09 | 9.9e-03 | 15 |
| Mycobacterium | Bacillus | 4.6e-04 | -39.78 | 9.9e-03 | 3 |
| Quinic acid | 3-Methyl-L-histidine | 4.6e-04 | -39.63 | 9.9e-03 | 4 |
| Jatrophihabitans | Asparagine | 4.6e-04 | -39.53 | 9.9e-03 | 3 |
| Pseudoflavitalea | Threonic acid hemicalcium salt | 4.5e-04 | -38.84 | 9.9e-03 | 2 |
| Abscisic acid | Fructose-6-phosphate | 4.5e-04 | -38.66 | 9.9e-03 | 4 |
| 3-Hydroxyanthranilic acid | Flavanomarein | 4.5e-04 | -38.56 | 9.9e-03 | 6 |
| Fructose-6-phosphate | Genistin | 4.5e-04 | -38.43 | 9.9e-03 | 3 |
| Devosia | GABA | 4.4e-04 | -37.80 | 9.9e-03 | 2 |
| Malic acid | Chr12_8480223 | 4.3e-04 | -36.52 | 9.9e-03 | 6 |
| Rhodanobacter | Ornithine | 4.1e-04 | -35.06 | 9.9e-03 | 7 |
| 2,3-Diaminopropionic acid | Orobol | 4.0e-04 | -34.61 | 9.9e-03 | 3 |
| Suberic acid | Daidzin | 4.0e-04 | -34.45 | 9.9e-03 | 13 |
| Ramlibacter | Daidzin | 3.9e-04 | -33.92 | 9.9e-03 | 10 |
| Ramlibacter | 2,3-Diaminopropionic acid | 3.8e-04 | -32.54 | 9.9e-03 | 4 |
| Succinic acid | Malic acid | 3.8e-04 | -32.43 | 2.0e-02 | 3 |
| Indole-3-carboxyaldehyde | 2'-Hydroxydaidzein | 3.6e-04 | -30.79 | 9.9e-03 | 15 |

**Table B3**: SHAP interaction edges with negative Z-scores ($Z \leq -30$). *weight* denotes the interaction difference $|\Delta|$ (Drought − Control), with two-sided $p$-values and bootstrap stability.

### Appendix C Implementation

#### C.1 Analysis

Statistical analyses and data visualizations were performed using a combination of R (version 4.4.1) and Python (version 3.10.9, Clang 14.0.6). Figures created in R primarily used the `ggplot2` package (version 3.5.1), while Python-based analyses—including SHAP value interpretation and XGBoost modeling—were conducted via the `reticulate` interface in RStudio. All the codes are available on GitHub: (https://github.com/Yoska393/ShapPlantMicro)

#### C.2 Hyperparameter tuning

We implemented two Random Forest (RF) models for phenotypic prediction using omics data. RF is a robust machine learning method that reduces overfitting through bootstrapping. It includes several hyperparameters, such as tree depth and the number of trees, which influence performance.

The first model used the `ranger` implementation (Wright and Ziegler, 2017) to predict biomass-related phenotypes based on the full omics dataset (8,757 predictors). Each response was modeled separately, and the mean performance was reported.

- `mtry` $\in \left\{ \left\lfloor \frac{\text{ncol}(X)}{6} \right\rfloor, \left\lfloor \frac{\text{ncol}(X)}{3} \right\rfloor, \left\lfloor \frac{\text{ncol}(X)}{1.5} \right\rfloor \right\}$: the number of variables randomly sampled as candidates at each split. Smaller values introduce more randomness and can help prevent overfitting.
- `min.node.size` $\in \{3, 5, 7\}$: the minimum number of observations in a terminal (leaf) node. Smaller values allow the model to grow deeper trees, potentially capturing more detail but increasing the risk of overfitting.
- `num.trees` $\in \{100, 500, 1000\}$: the total number of trees in the forest. More trees generally improve stability and performance, but increase computation time.

The second model used XGBoost (Chen and Guestrin, 2016) for predicting a representative dry weight phenotype using a reduced feature set (2,729 features). The variable inputs were selected based on feature importance scores from the first model. The XGBoost hyperparameter grid included:

- `learning_rate` $\in \{0.1, 0.3, 0.5\}$: the step size shrinkage used in each boosting step. Lower values make the model more robust but require more boosting rounds.
- `n_estimators` $\in \{50, 100, 500\}$: the number of boosting rounds (trees). More rounds can improve performance, but may lead to overfitting if not regularized.
- `max_depth` $\in \{3, 6, 9\}$: the maximum depth of each tree. Shallower trees are more regularized and less prone to overfitting, while deeper trees can model complex patterns.

Both models were evaluated using 10-fold cross-validation under control and drought conditions, with mean squared error (MSE) as the evaluation metric. To ensure a fair comparison, we aimed to select identical hyperparameters for both conditions whenever possible.

The grid search results, summarized in Fig. B3, show minimal differences in performance across all hyperparameters. RF has proven robust under all tested configurations, likely because of its bootstrapping strategy. We object to selecting the same hyperparameter set, which is stable under both control and drought conditions. Considering overall performance, overfitting risks, and computational efficiency, we selected the following default configurations:

- **Ranger**: `mtry` = $\left\lfloor \frac{\text{ncol}(X)}{3} \right\rfloor$, `min.node.size` = 5, `num.trees` = 500
- **XGBoost**: `learning_rate` = 0.3, `n_estimators` = 100, `max_depth` = 6

#### C.3 Stability-Based Filtering of Differential Interactions

Under the present modeling and feature-selection setting, SHAP interaction matrices are extremely sparse. In a single bootstrap replicate, the observed edge density was approximately 0.015% (5,563 edges out of 3,722,356 possible feature pairs).

As a rough baseline, the appearance of a specific interaction edge in a given bootstrap replicate can be treated as a rare event with probability $p \approx 1.5 \times 10^{-4}$. Under this assumption, the number of times an edge is observed across $B_{\text{boot}} = 100$ bootstrap replicates can be approximated by a binomial distribution:

$$X \sim \text{Binomial}(100, p).$$

Although it may appear counterintuitive, this null model implies that repeated detection of the same edge, even in a small number of bootstrap replicates, is already highly unlikely to occur by chance alone. Specifically, the probabilities that a given edge appears in at least two, three, or four Bootstrap replicates are given by

$$\Pr(X \geq 2) = 1 - \sum_{k=0}^{1} \binom{100}{k} p^k (1-p)^{100-k} \approx 1.1 \times 10^{-4},$$

$$\Pr(X \geq 3) = 1 - \sum_{k=0}^{2} \binom{100}{k} p^k (1-p)^{100-k} \approx 5.4 \times 10^{-7},$$

and

$$\Pr(X \geq 4) = 1 - \sum_{k=0}^{3} \binom{100}{k} p^k (1-p)^{100-k} \approx 2.0 \times 10^{-9}.$$

Thus, observing the same interaction in only 2–4 out of 100 bootstrap replicates already provides non-trivial statistical evidence against a purely random, sparsity-driven null model.

Based on this reasoning, we adopted a conservative stability-based filtering criterion requiring an edge to be observed in at least two bootstrap replicates ($n_{ij}^{\text{obs}} \geq 2$). This threshold effectively removes isolated, non-reproducible edges that are likely to arise from resampling noise, while retaining interactions that are consistently recovered across bootstrap samples.
